# Morpho-molecular features of Epithelial Mesenchymal Transition associate with clinical outcome in patients with rectal cancer

**DOI:** 10.1101/2024.11.07.622481

**Authors:** Mauro Gwerder, Cansaran Saygili Demir, Hannah L. Williams, Alessandro Lugli, Cristina Graham Martinez, Joanna Kowal, Amjad Khan, Philipp Kirchner, Thibaud Koessler, Martin D. Berger, Martin Weigert, Inti Zlobec

**Affiliations:** Institute of Tissue Medicine & Pathology, University of Bern, Bern, Switzerland; Graduate School for Cellular and Biomedical Sciences, University of Bern, Bern, Switzerland; Lunaphore Technologies, Tolochenaz, Switzerland; Department of Oncology, Geneva University Hospitals, Geneva, Switzerland; Department of Medical Oncology, Inselspital, Bern University Hospital, University of Bern, Bern, Switzerland; Center for Scalable Data Analytics and AI (ScaDS.AI), Dresden/Leipzig, Germany; Faculty of Computer Science, TU Dresden, Dresden, Germany; Institute of Bioengineering, School of Life Sciences, École Polytechnique Fédérale de Lausanne (EPFL), Lausanne, Switzerland

**Keywords:** Rectal cancer, tumor budding, epithelial-mesenchymal transition, neoadjuvant chemoradiotherapy, hyperplex immunofluorescence

## Abstract

In rectal cancer, where part of the patients undergoes chemoradiotherapy, there is a need for improved pretreatment biomarkers applicable to biopsies. Tumor budding (TB) is a biomarker used in colon cancer, and due to its link to epithelial-mesenchymal transition (EMT), is hypothesized to be a potential marker for therapy resistance. Assessment of the utility of tumor buds in rectal biopsies is challenging due to their rarity. As EMT-related processes are also seen in other morphological features beyond tumor buds, we investigated EMT in tumor tissue including morphological features such as tumor *cluster size* and fibril-like structures. To do so, we leveraged a cohort of colon cancer whole-slide images and another cohort consisting of rectal cancer biopsies, visualized using hyperplex immunofluorescence to identify tumor and EMT-associated proteins. We built a custom image analysis pipeline to detect and segment tumor buds and other morphological features and correlated them with molecular expression intensities. We found strong correlations of EMT up-regulation and morphological transition states, both at the invasive margin and the tumor center. We furthermore observed a link between morpho-molecular transitions and histological growth patterns, which in turn can inform novel biomarkers. Finally, quantification of these morpho-molecular transition states in rectal biopsies showed their impact on survival after neoadjuvant chemoradiotherapy.

## 1 Introduction

Colorectal cancer (CRC) is the third most prevalent cancer type in Switzerland and worldwide, of which roughly 40% are of rectal origin [1]. Although rectal and colon cancers are molecularly comparable, treatment strategies differ strongly due to anatomical differences [2]. Total neo-adjuvant therapy (TNT) is standard of care for locally advanced rectal cancer (LARC) [3], [4], [5] to facilitate subsequent tumor resection [6] and in some cases guarantee organ preservation [5]. Future treatment strategies could be refined to incorporate two needs: Firstly, more patients responding well to treatment are suited to a watch-and-wait strategy, where no surgery is needed [7], [8]. Secondly, pre-treatment assessment of biopsies could guide patient selection for treatment regimens [9]. To make this paradigm a reality, novel biomarkers are needed to improve prediction of response from pre-treatment specimens such as biopsies. Although histopathological assessment of biopsies is routinely conducted, it acts only as a confirmation of cancer incidence rather than informing treatment decision. In the context of post-treatment resections, pathologists report post-treatment staging (ypTNM) and tumor regression grading (TRG) [10], [11], [12], [13]. The prognostic value of TRG is unclear, due to high inter-observer variability and definition differences between grading schemes [14]. New pre- and post-treatment response scoring methods are promising [15], [16], [17], however, further studies are needed to uncover underlying biological mechanisms of these methods.

A potential avenue for novel response biomarkers is tumor budding (TB), a widely accepted and reported hallmark of aggressive, infiltrative colon cancer [17], [18]. TB is used as a biomarker for worse prognosis, specifically for metastatic progression of the disease [19]. Defined as tumor clusters consisting of four or less cells, pathologists routinely assess TB in surgically treated colorectal cancer [20], where tumor buds are counted at the invasive margin of the primary tumor, according to guidelines given by the International Tumor Budding Consensus Conference (ITBCC) [20]. In contrast, TB is not an established biomarker in locally advanced rectal cancer (LARC) for two reasons: First, the standard of care for LARC patients is to undergo neoadjuvant long course chemoradiotherapy (nCRT) followed by surgery [21], [22]. Thus, assessing TB in post-treatment resections is unreliable, and therefore not recommended [20], as remains of apoptotic tumor fragments can be mistaken for tumor buds. Second, TB is also not assessed routinely in biopsies, as they are taken from the tumor center. According to ITBCC recommendations, pathologists only score TB at the invasive margin [20]. This scoring method does not translate to biopsies, as they are routinely taken from the tumor center. Although intratumoral budding (ITB) does exist and is also associated with worse prognosis [20], it is a rarer feature than peritumoral budding (PTB).

Despite its rarity, ITB remains a scenario for novel biomarkers, as hypothesized by the ITBCC due to its reported prognostic value in colon and rectal cancers [23], [24], [25]. Specifically, Rogers et al. [24] showed the association of treatment response and ITB. Such results can be explained molecularly: tumor buds are hypothesized to undergo epithelial-mesenchymal transition (EMT). Cells undergoing EMT lose their adhesiveness to other epithelial cells and up-regulate a variety of processes including motility, making it easier for them to move through stromal tissue [26] to ultimately invade surrounding vessels and lymphatics [27]. Tumor buds also display reduced proliferation during that process [27]. Specifically, tumor buds are hypothesized to enter a partial EMT state [28], where epithelial adhesive markers such as E-cadherin, Catenin beta-1 and Ep-CAM are down-regulated at the cell membrane. In CRC, tumor buds also show down-regulation of CDX2, a homeobox transcription factor for intestinal differentiation [29]. In contrast, tumor buds have been shown to up-regulate mesenchymal markers such as Vimentin [30] and transcription factors such as ZEB1 [31] and nuclear Catenin beta-1 [32]. EMT itself is linked to therapy resistance [33], [34], although this behavior is hard to prove in a translational setting.

In addition to TB, other related morphological biomarkers have been suggested for colorectal cancer. For example, poorly differentiated clusters (PDCs) are defined as tumor clusters larger than four cells without an apparent lumen [35]. Furthermore, infiltrative growth patterns at the invasive margin, often accompanied with fibril-like growth, have previously been associated with worse prognosis [36]. Interestingly, there is evidence of a connection to EMT for both features. Tumor *cluster size* and EMT-related marker expression is negatively correlated as shown by Lin et al. [37], indicating that EMT is already detectable in larger clusters than tumor buds, although to a lesser degree. Therefore, PDCs could represent a conduit between tumor mass and tumor buds. Furthermore, fibril-like structures extending away from the main tumor body are linked to the same EMT-related molecular changes [38]. In fact, TB at the end of fibril-like structures was reported by Lin et al. [37], suggesting tumor buds “sprouting off” of fibrils. However, to the best of our knowledge, no study so far has compared the transitional states of EMT between fibril-like tumor extensions, TB and PDCs.

The underlying molecular changes that drive these morphological features are complex, suggesting that the simultaneous visualization of multiple proteins is needed to sufficiently characterize them [39]. Hyperplex immunofluorescence [40], and more specifically sequential immunofluorescencë[41] has recently emerged as a powerful method to visualize more than 100 proteins on the same histological section. Using a cyclical approach, primary and secondary antibodies are administered, imaged, and eluted from the tissue section. Through the visualization of many markers within the same section of tissue, this method enables a deeper understanding of the molecular landscape of the tissue while retaining spatial architecture. This in turn will inform the construction of smaller panels to facilitate clinical implementation of derived biomarkers of response [39].

In this study, we hypothesize that fibril-like structures, PDCs, and tumor buds are all part of the same continuous morphological process, representing EMT. We propose further that this phenotypical transition together with morphological features is not only observable in the context of rectal cancer biopsies but is also prognostic of patient survival. To do so, we designed a sequential immunofluorescence panel consisting of 31 protein targets, focused on capturing epithelial cell marker expression together with histomorphological features. We applied the established panel to a small set of colon cancer cases, acting as a discovery and pipeline development cohort. After confirming the viability of the custom computational pipeline by replicating previous results [29], [37], we applied the workflow to a large rectal cancer biopsy cohort, proving the clinical relevance of morpho-molecular transition states in a pre-treatment context.

## 2 Results

To investigate morpho-molecular transition states for novel biomarker discovery in rectal cancer biopsies, two colorectal cancer cohorts were selected. Five whole-slide images (WSI) were acquired from a cohort of colon cancer resections. These images were utilized for seqIF™ panel establishment and the validation of the custom pipeline for *cluster size* analysis (described in detail in 4.3). A second cohort consisting of 160 rectal cancer biopsies with follow-up data was subsequently leveraged to apply the established pipeline and find relations to patient outcome. This second cohort is a tissue micro array (TMA), where each biopsy is represented by a sampled tissue core (∅= 1*mm*; see 4.1).

As a first step in the custom seqIF™ analysis pipeline, the tumor tissue is segmented into tumor clusters. A tumor cluster is defined as a population of cells that are adhered to each other and are fully surrounded by stromal tissue. Afterwards, tumor clusters are classified as tumor buds and non-budding tumor clusters (defined as tumor clusters consisting of more than 4 cells; Fig. 1a).

**Fig. 1.**
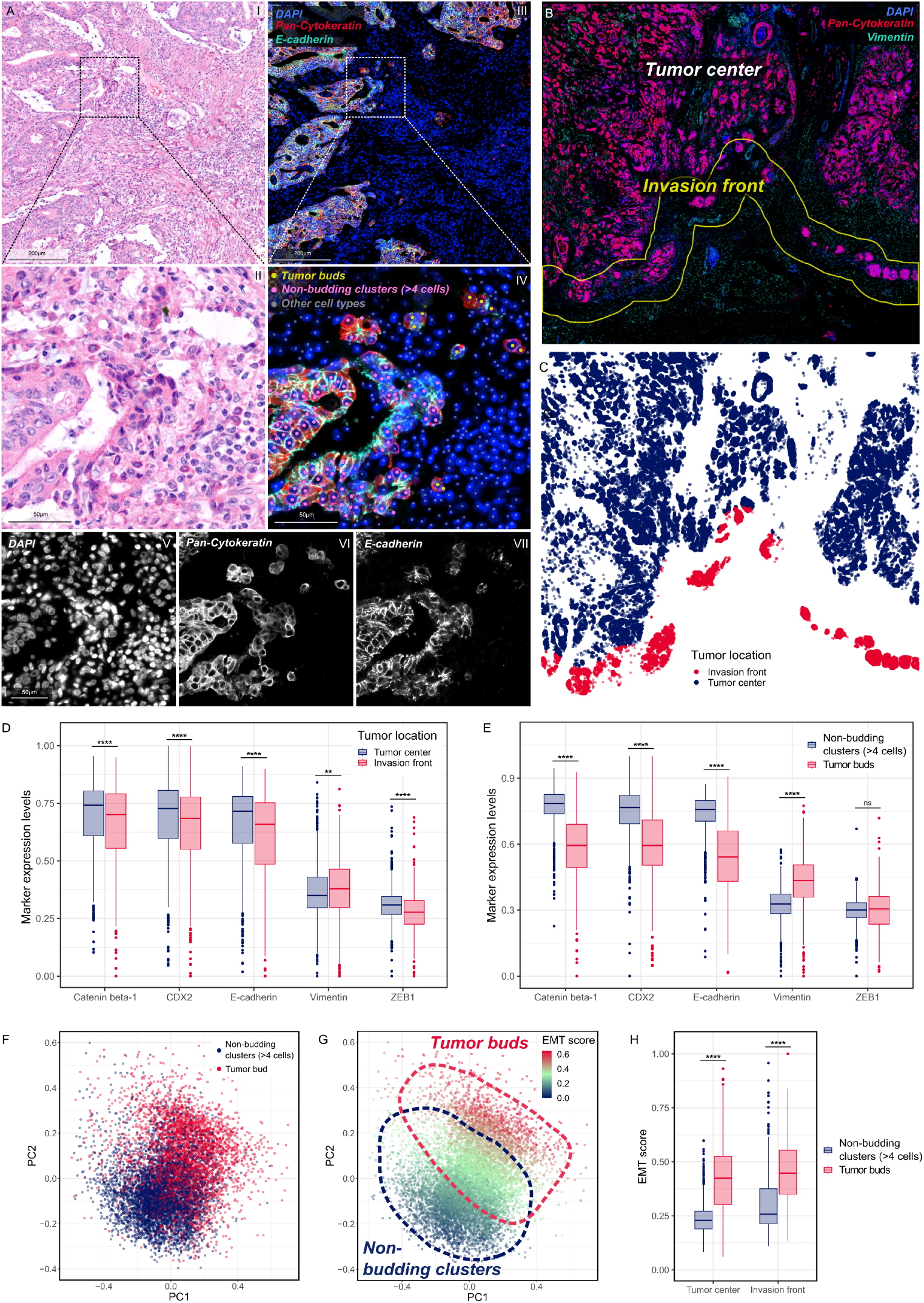
**(A)** H&E images (I&II) and seqIF™ images (III & IV) of the same tumor area, taken from image 2 of the colon cancer cohort. Classification of each cell is given with colored centroids (IV). Single channel images are shown in (V-VII). **(B)** Manual annotations of tumor center and invasion front (yellow), visualized on image 4 of the colon cancer cohort. **(C)** Cell centroids are labeled based on their location in the “tumor center” or the “invasive front”. **(D)** Differential mean marker expressions in both tumor locations “tumor center” and “invasion front”. Each data-point corresponds to a tumor cluster. **(E)** Differential marker expression between the tumor cluster classes “tumor buds” and “non-budding clusters”. Bulked non-budding clusters are compared with bulked tumor bud clusters of the same image tile. **(F)** PCA visualization of each cluster’s mean expression, with cluster classes as their labels. **(G)** PCA visualization with *EMT score* as their labels. **(H)** Comparison of *EMT score* values between tumor cluster classes “tumor buds” and “non-budding clusters”, evaluated in both tumor locations.

### 2.1 Exploring the influence of location on tumor budding

To potentially understand the driving forces of EMT, it is important to first explore differential marker expression in a spatial context. Therefore, we have compared the invasion front with the tumor center in terms of EMT up-regulation. Before grouping tumor cells into clusters, regions of tumor tissue from five WSIs were manually annotated as either “tumor center” or “invasion front”. The invasion front was defined as the area surrounding the deepest cancer invasion, with a radius of 500*µm* (Fig. 1b) [42]. Each tumor cell was assigned a label according to their spatial location (Fig. 1c). We subsequently compared marker expression intensities for five EMT-related protein markers (Catenin beta-1, E-cadherin, CDX2, Vimentin, ZEB1), with potential expression in epithelial cells (Fig. 1d). A more detailed description of these markers can be found in table 1. A description of the full panel, including markers that can capture immune invasion, cell status as well as cancer-associated fibroblasts, can be found in table 2.

**Table 1.** Description of proteins that are potentially expressed in epithelial and TB cells.

| Protein name | Description of relevance in TB and EMT | Sources |
| --- | --- | --- |
| Catenin beta-1 | Down-regulated as adhesion molecules at the cell membrane in EMT | [63], [64], [65], [66], [32] |
| CDX2 | Intestine-differentiation transcription factor often lost in CRC at the invasion front | [67], [29] |
| E-cadherin | Down-regulated as adhesion molecules at the cell membrane in EMT; can also act as a transcription factor and is reported to move to the cell nucleus during Wnt-pathway deregulation and EMT. | [64], [65], [66], [68] |
| CD47 | CD47 has been linked to immune-escape. There is no direct link of CD47 to EMT. | [69] |
| ZEB1 | transcription factor in the EMT-associated Wnt-pathway | [31] |
| Ep-CAM | Epithelial marker shown to be up-regulated in colorectal cancer and down-regulated in EMT. | [57], [70], [71] |
| Lamin-B1 | Linked to EMT in non-small cell lung cancer; more broadly associated to cellular senescence | [59], [60] |
| Ki-67 | Proliferation marker; cells undergoing EMT decrease proliferation | [27], [72], [73] |
| Caspase-3 | Small cell clusters expressing Caspase-3 are not likely to be viable nor relevant for metastatic progression | [58] |
| Vimentin | Vimentin-expression in tumor buds is indicative of a partial EMT state | [30] |

**Table 2.**
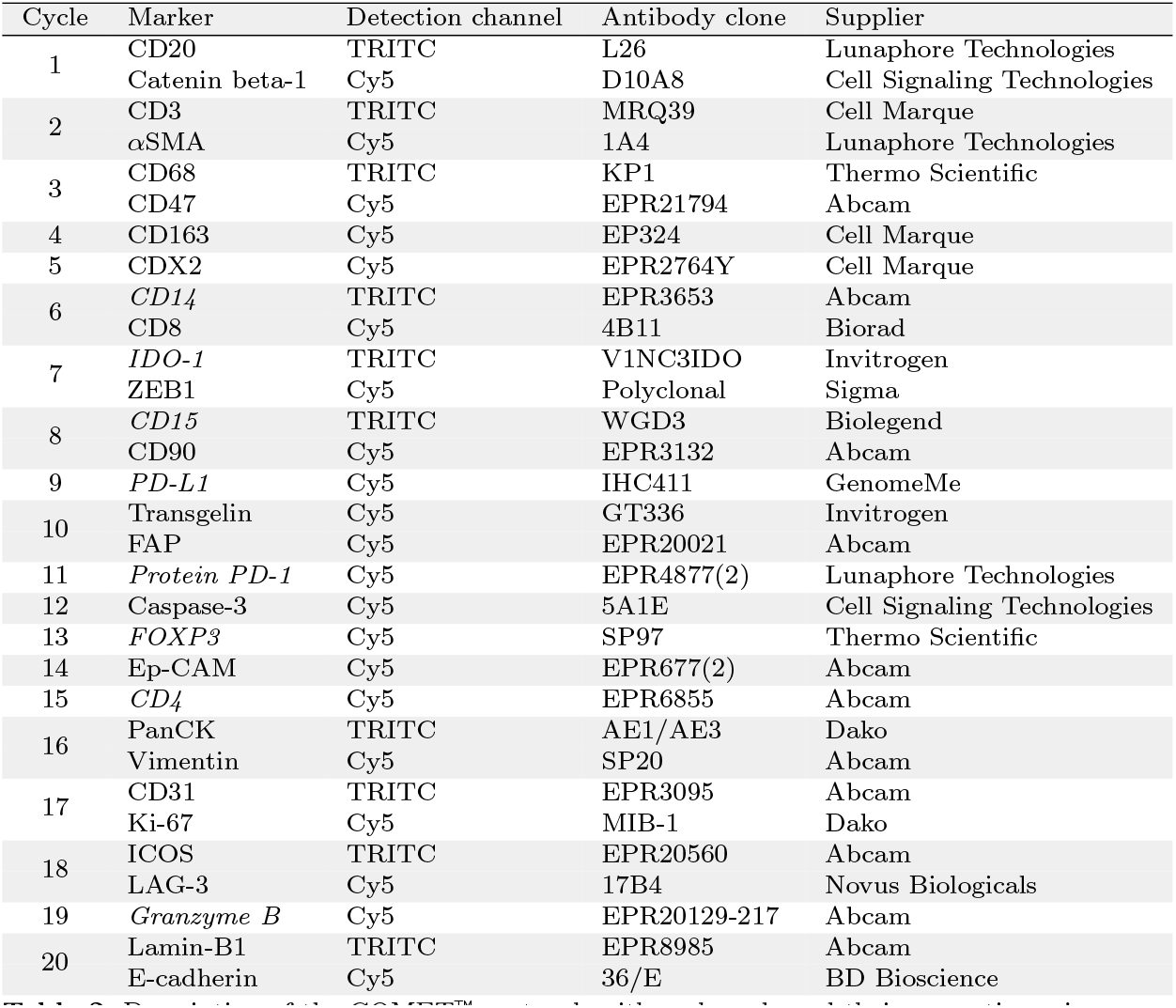
Description of the COMET™ protocol, with each cycle and their respective primary antibodies listed. Markers in cursive have been applied only to the rectal biopsy cohort, making up the final 31-plex panel.

We observed a significant down-regulation of membrane-bound Catenin beta-1, CDX2 and E-cadherin at the invasion front compared to the tumor center (*p* < 0.0001, with effect sizes cohen’s d of 0.24, 0.31 & 0.25 respectively; Fig. 1d). However, as TB is more common at the invasion front, this difference is likely driven by the number of tumor buds and smaller tumor clusters (0.8% of all tumor cells in the tumor center are budding cells, compared to 1.4% at the invasion front). As average expression levels of markers ZEB1 and Vimentin are below a normalized value of 0.5, these marker expression differences were considered background noise for the purposes of this study (see 4.3.3). Catenin beta-1 expression was also down-regulated in the nuclear compartment (supp. Fig. 1), contrary to findings showing Catenin beta-1 moving to the nucleus to act as a transcription factor [32].

Subsequently, the custom pipeline detected tumor clusters and classified them as tumor buds or non-budding tumor clusters (Defined as any tumor cluster containing more than 4 cells; see 4.4). When comparing tumor buds to non-budding tumor clusters (Fig. 1e), membrane-bound Catenin beta-1, CDX2, E-cadherin were down-regulated (*p* < 0.0001, with effect sizes cohen’s d = 1.27, 1.06, 1.57 & 0.97 respectively). Due to the overall low expression, Vimentin up-regulation in tumor buds (effect size cohen’s d = 0.98) is most likely caused by signal spillover from neighbouring mesenchymal cells. Qualitative investigation of Vimentin expression in TB confirmed this hypothesis. Finally, ZEB1 up-regulation was not observed in TB. To summarize, we showed down-regulation of relevant epithelial markers associated with EMT in at the invasive front and specifically in TB. However, Vimentin, ZEB1 and nuclear Catenin beta-1 up-regulation, as previously reported in TB [30], [31], [32], could not be confirmed. To resolve more global phenotypic differences between tumor buds and non-budding tumor clusters, the principal components of each cluster’s expression pattern were analyzed (Fig. 1f). The resulting PCA showed that both groups show substantial overlap, suggesting that the heterogeneity of the tumor cell clusters could be represented better using a continuum rather than a binary state. To represent this transition in a quantitative manner we derived an *EMT score* by calculating the average expression of three EMT-related, strongly down-regulated markers (Catenin beta-1, CDX2 & E-cadherin) per cell (see 4.5). This EMT score calculation is inspired by similar approaches in transcriptomic research [43], [44]. This score gradually increases from non-bud clusters to tumor buds (Fig. 1g). A visualization of the EMT score on tumor tissue is given in supplementary figure 3. As biopsies are taken from the tumor center rather than the invasion front [20], we set to investigate differences between TB and non-TB clusters in both tumor locations. Indeed, *EMT scores* in TB and non-TB clusters are characterized by up-regulation of EMT regardless of their spatial context. When comparing *EMT scores* between TBs and non-TB clusters, there were higher levels in both tumor locations (*p* < 0.0001, with effect sizes cohen’s d = 1.58 within the tumor center & 1.07 at the invasive margin; Fig. 1h). These findings suggest that EMT-related changes of TB marker expression are independent of tumor location, and could therefore be detectable in intratumoral buds in rectal biopsies as well.

### 2.2 Budding in rectal cancer biopsies: A glimpse into intra-tumoral EMT activity

To find novel EMT biomarkers for rectal cancer treatment outcome, it is necessary to confirm that TB in rectal cancer biopsies demonstrate similar evidence of EMT-related processes as well. After validating the established analysis pipeline on WSI, the same pipeline was applied to the rectal cancer biopsy cohort, comprising 160 patients (detailed description in 4.1). New annotations for this second cohort were made by a trained scientist (see 4.4). An extended version of the original panel was employed, which elaborated on epithelial and cell status markers (Ep-CAM, Ki-67, Caspase-3, Lamin-B1; see 4.2). A more detailed description can be found in table 1. To provide an overview of the epithelial markers, co-expressions of available epithelial and cell-status markers were calculated (Fig. 2a). Highest correlation values are found between E-cadherin, Catenin beta-1 and Ep-CAM. Furthermore, there is a correlation between the nuclear markers CDX2 and Lamin-B1.

**Fig. 2.**
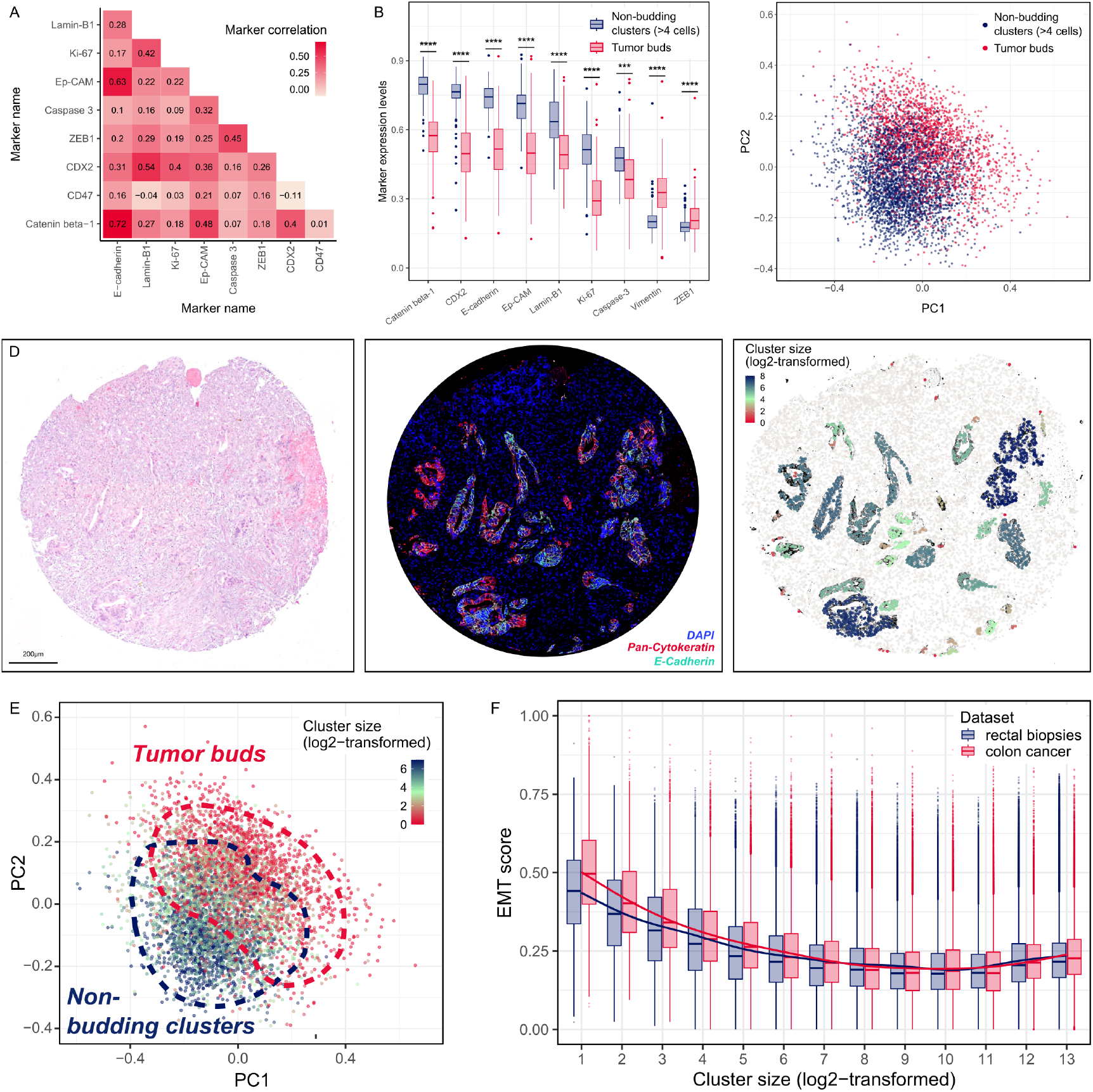
**(A)** Marker co-expression correlations, calculated on the rectal biopsy cohort. **(B)** Differential marker expression between the tumor cluster classes “tumor buds” and “non-budding clusters” on the rectal biopsy cohort. **(C)** PCA visualization of each cluster’s mean expression, with cluster classes as their labels, calculated on the rectal biopsy cohort. **(D)** Visualization of the log2-transformed *cluster size* feature. In addition, matching H&E and IF images of the same TMA core are shown. **(E)** PCA visualization of each cluster’s mean expression, with *cluster size* (log2-transformed) as their label. **(F)** Correlation of *EMT score* and *cluster size* (log2-transformed) for both cohorts.

Of the 160 patient cores, 79 showed intratumoral budding. Down-regulation of markers Catenin beta-1, CDX2 and E-cadherin was detected in tumor buds, showing that EMT-related processes can be identified in biopsies (Fig. 2b; *p* < 0.0001, with effect sizes cohen’s d 2.5, 2.34 and 2.15 for markers Catenin beta-1, CDX2 and E-cadherin respectively). Additionally, we found significant down-regulation of Ki-67, Ep-CAM, Lamin-B1 and Caspase-3 in tumor buds (*p* < 0.0001, with effect sizes cohen’s d = 1.62, 1.78, 1.15 and 0.80).

To summarize, differential marker expression in rectal biopsies confirmed the findings on colon cancer resections. Additionally, markers Ki-67, Ep-CAM, Caspase-3 and Lamin-B1 were down-regulated in intratumoral TB in rectal cancer. Furthermore, we identified gradual phenotypic changes between tumor buds and non-budding clusters (Fig. 2c) as previously identified in the colon cancer WSIs. Compared to the colon cancer cohort, even weaker separation between both groups was observed. Nevertheless, we were able to confirm that TB is also related to EMT in rectal biopsies and is a potential avenue for novel biomarkers.

### 2.3 Beyond tumor buds: The impact of tumor cluster size

Although the morphomolecular representation of EMT is detectable in rectal cancer biopsies, the binary classification of tumor buds and non-budding clusters does not capture the transition seen in both cohorts to its full extent. This is especially an issue in consideration of the rarity of intratumoral budding (ITB). We therefore explored the correlation of the continuous variable *cluster size* with EMT. To assess *cluster size*, the number of tumor cells within each uniquely segmented tumor area was counted. This value was subsequently assigned to each cell within the cluster (Fig. 2d). As expected, *cluster size* is resolving the molecular transition more finely than the previous binary classification Fig. 2e). Additionally, a negative correlation of EMT score and *cluster size* in both cohorts was observed, with a stronger effect in the colon cancer cohort (Spearman *ρ* = −0.35 as compared to *ρ* = −0.32 in the colon cancer cohort, Fig. 2f). This difference between the cohorts can be explained by the substantial punch artifacts and disconnected tumor clusters present in the TMA data set. Furthermore, the correlation seems to trail off at around a *log*(2)-transformed *cluster size* of 7.0 (corresponding to *cluster sizes* of 128 cells). These results are suggestive of a prognostic value for tumor *cluster size* beyond the threshold-based definition of tumor buds. Investigating individual epithelial markers and their relation to *cluster size*, we find correlations with markers Catenin beta-1, E-cadherin, and CDX2 (supp. Fig. 4-5 and Fig. 4h). We were not able to replicate the correlation with Ep-CAM and Lamin-B1 in the rectal biopsy cohort (supp. Fig. 6-8). Expression levels of these markers seem to be influenced more strongly by inter-patient differences rather than *cluster size*. Ki-67 showed marker level upregulation only in very large clusters (*cluster size¿* 8000; supp. Fig. 9) However, a limited number of WSI samples was analyzed with the full panel. Therefore, these findings will need to be confirmed with further investigation. Overall, treating *cluster size* as a continuous feature correlates well with EMT score and enables us to bypass the limitation of ITB rarity.

### 2.4 Deciphering tumor clusters: EMT found in fibril-like structures

Continuing the inspection of morphological features, EMT-related changes could be detected even within one tumor cluster: fibril-like structures are implicated to also correlate with EMT [37]. We qualitatively observed EMT up-regulation in fibril-like structures in the colon cohort. To quantify this, we determined an *interaction score*, where unbiased metrics were used to quantitatively assess fibril-like infiltrative growth patterns. Around each cell within a radius of 30*µm*, the number of cancer-associated fibroblasts (CAF; cell-typing is described in 4.3.3) is counted, quantifying the degree of infiltrative growth (*stromal interaction*; Fig. 3a II). To also capture the opposite growth pattern which results in bulky, compact cancer growth, the number of other epithelial cells within 30*µm* of each tumor cell was quantified (*Epithelial interaction*; Fig. 3a III). We then subtracted *epithelial interaction* values from *stromal interaction values* to create the *interaction score* (Fig. 3a IV). In doing so, we create a score per cell in which positive values signify stronger stromal than epithelial interaction. This corresponds to infiltrative growth and fibril-like structures, whereas negative values show that the epithelial interaction predominates, indicating compact growth.

**Fig. 3.**
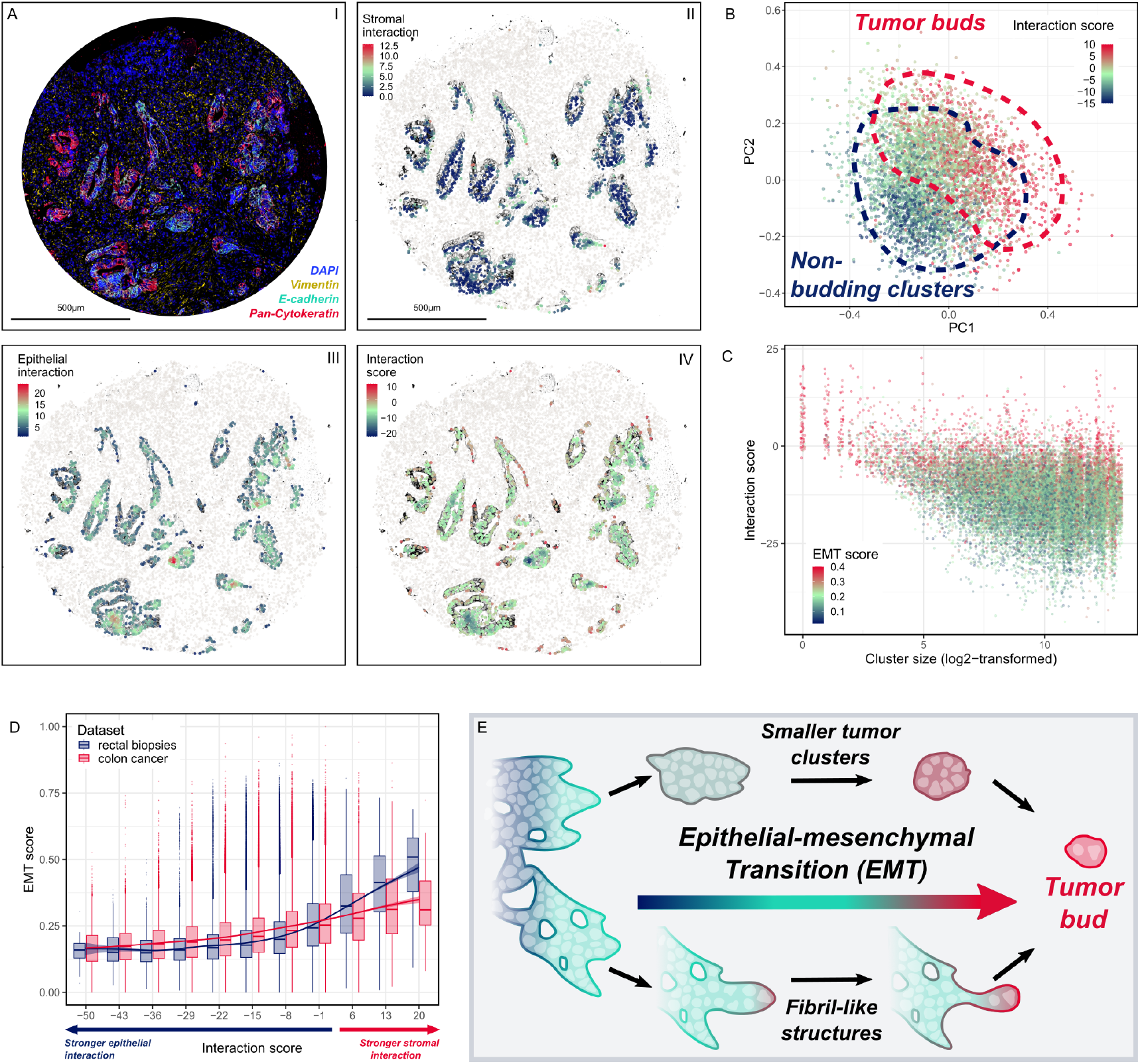
**(A)** Visualization of each component of the *interaction score*. **(B)** PCA visualization of each cluster’s mean expression, with *interaction score* as their labels. **(C)** Scatter plot showing the correlation of *cluster size* and *cluster size* (log2-transformed) in individual cells. Cells with high EMT score are also found in large tumor clusters and this is correlated with a high *interaction score*. This figure highlights that *EMT score* is also upregulated in large tumor clusters, as long as the *interaction score* of an individual cell is high. **(D)** Correlation of *EMT score* and *interaction score* in individual cells for both cohorts. **(E)** Graphical demonstration of how epithelialmesenchymal transition and morphological features can be related. Tumor buds are hypothesized to be at the end of a gradual change of morpho-molecular feature expression.

Inspecting the principal components of each tumor cluster, it is apparent that TB shows mostly positive values for *interaction scores*. As tumor buds are embedded within the stromal compartment, this finding is expected. Interestingly, some non-budding clusters show positive *interaction score* as well, suggesting that those specific clusters show infiltrative growth patterns (Fig. 3b). Moreover, when analyzing individual cells, a high *interaction score* coincided with EMT up-regulation (Fig. 3c). This was observed across all *cluster sizes*. Since the *interaction score* is higher at the edge of a cluster and highest in fibril-like structures, this indicates that up-regulation of EMT is a defining feature of tumor cells in intense contact with tumor stroma. This shows the added value of per cell evaluation of the *interaction score*, uncovering EMT-related changes in large tumor clusters as well. Finally, we show EMT up-regulation in both cohorts for positive *interaction score* values (Fig. 3d), suggesting that fibril-like extensions into the stroma is correlated with EMT up-regulation. The weaker effect in the colon cancer cohort can be explained by patient heterogeneity, further discussed in 2.5. Akin to results in 2.3, down-regulation of CDX2, Catenin beta-1 and E-cadherin is negatively correlated with positive *interaction score* values (Fig. 4i and supp. Fig. 12-13), suggesting that stronger stromal interaction correlates with EMT activation. We therefore show that fibril-like structures are another morphological feature associated with EMT, potentially representing a preliminary stage of TB (Fig. 3e).

### 2.5 Infiltrative growth patterns: Aligning molecular heterogeneity with histology

Both morphological parameters (*cluster size* and *interaction score*) showed considerable heterogeneity. To assess the inter-patient heterogeneity of the colon cancer cohort, we inspected the invasion fronts on matching H&E-stained sequential cuts. Clear differences in tumor histology between images became apparent: Image 1 shows medullary growth patterns [45] (Fig. 4a). Image 2 displays a compact and uniform invasion front (defined as “pushing tumor border configuration” by Jass et al. [46]), while Images 3-5 are identified by an irregular invasion front, with frequently visible fibrils and TBs (defined as “infiltrative tumor border configuration” by Jass et al. [46]). Interestingly, when EMT scores in TB were compared between images, we observed EMT up-regulation in infiltratively growing cancers (Fig. 4b). More strikingly, EMT up-regulation scales more strongly with *cluster size* (Fig. 4c) and *interaction score* (Fig. 4d) in the case of infiltrative growth. This means that active EMT processes can be captured by quantifying morphological changes of the tumor, such as invasive growth patterns.

**Fig. 4.**
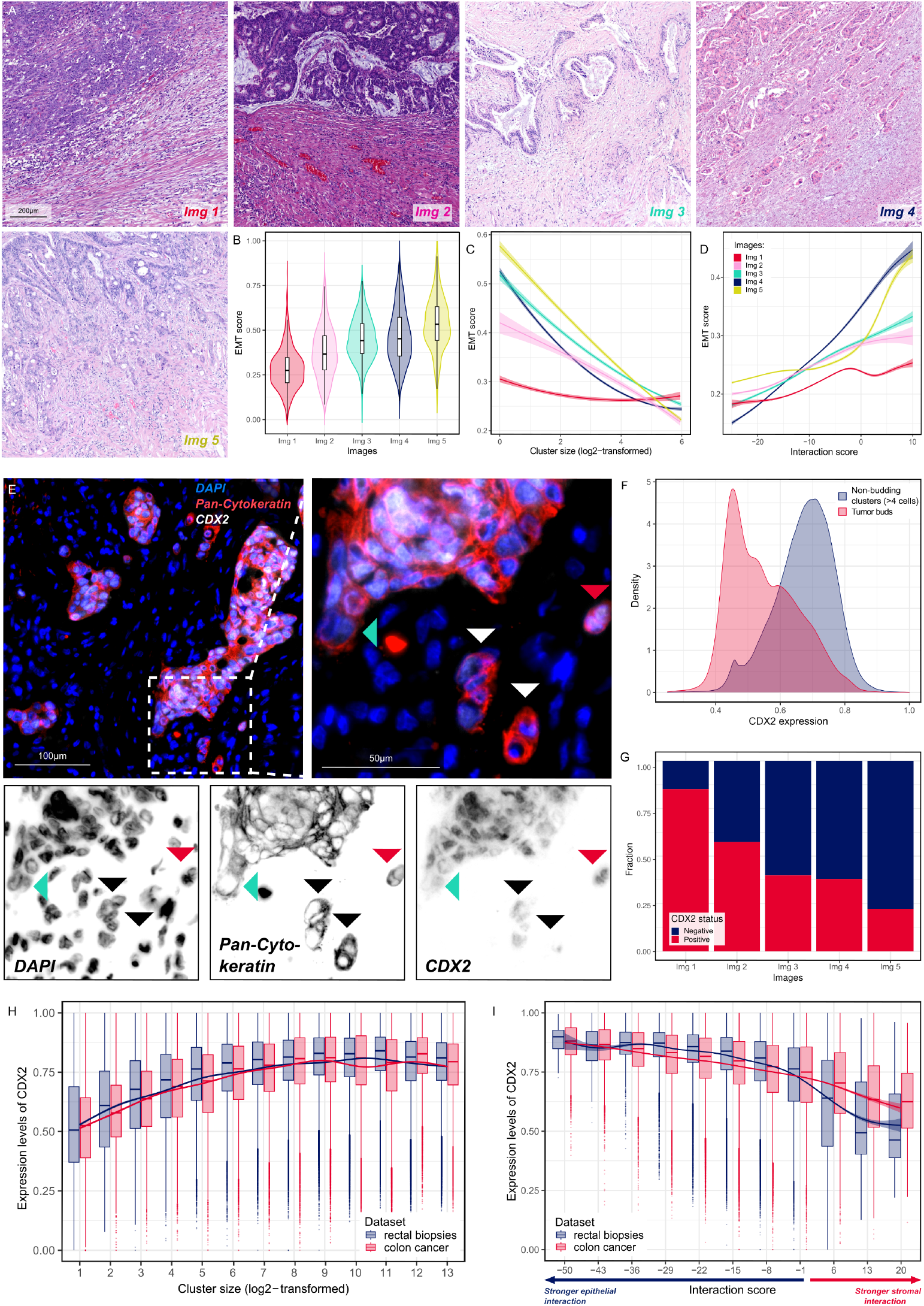
**(A)** Representative H&E images of the invasive margins in the colon cancer WSI. Image 1 shows medullary growth. Image 2 shows pushing border growth patterns. Images 3-5 are growing in an infiltrative manner. **(B)** Mean *EMT scores* across all tumor bud cells per whole slide image. **(C)** Correlation of *cluster size* and *EMT score*, given for each tumor cluster and each whole slide image. **(D)** Correlation of *interaction score* and *EMT score*, given for each tumor cell and each whole slide image. **(E)** The variability CDX2 expression in infiltrative tissue. Shown are representative examples of CDX2-negative tumor buds (white/black arrows), a fibril-like structure with CDX2 negativity (teal arrow) as well as a CDX2-positive tumor bud. **(F)** CDX2 expression densities of tumor buds and non-budding clusters. **(G)** Fraction of CDX2-positive (red) and CDX2-negative (dark blue) tumor buds per whole slide image. **(H)** Correlation of CDX2 expression and *cluster size*for individual cells in both cohorts. **(I)** Correlation of CDX2 expression and *interaction score* for individual cells in both cohorts.

Investigating the impact of single markers on their impact on tumor growth patterns, we found interesting patterns in the expression patterns of CDX2 (Fig. 4e). CDX2 is considerably downregulated in most tumor buds (black arrows) as well as in fibril-like structures (teal arrow). However, in comparison to other markers, there are also tumor buds showing clear CDX2 positivity (red arrow). Indeed, we observed a subpopulation of tumor buds expressing CDX2, and vice versa, tumor cells within non-budding tumor clusters with loss of CDX2 expression (Fig. 4f). This heterogeneity of CDX2 expression can also be explained by differences in morphology: Infiltrative-growing tumors show a lower fraction of CDX2-positive tumor buds (Fig. 4g). When trying to confirm this result in the biopsy samples, the number of confidently classified tumor buds in the rectal biopsy cohort remains very small, due to the high amount of artifacts and small tissue area. Introducing the *interaction score* as a filtering step (see 4.5) was not enough to exclude various artifacts such as ruptured glands [47]. To bypass this limitation, instead of solely focusing on TB, CDX2 expression and its correlation to *cluster size* (Fig. 4h) and *interaction score* (Fig. 4i) were investigated as well. These results are comparable to previous findings with *EMT score* (Fig. 2f & 3d).

Overall, we showed that well-known histological patterns (infiltrative vs. pushing border [36]) reveal active EMT processes. It becomes apparent that interplay of morphological and molecular features are both needed to reliably quantify infiltrative processes in colorectal cancers. However, these findings are limited to the WSI data, as such large scale histological patterns are only valid to be reported by pathologists in resections without neoadjuvant treatment. Nevertheless, correlation of local morphological and molecular features can also be assessed in rectal biopsies, potentially giving an estimate of the infiltrative potential of each tumor.

### 2.6 Leveraging morpho-molecular heterogeneity to predict patient survival

As a final step, the clinical relevance of identified molecular and morphological features in rectal cancer biopsies was assessed. The prognostic value of tumor regression grading is strongly debated [14], so we decided to focus on survival endpoints. Specifically, the potential to predict overall survival (OS) and disease-free survival (DFS) using morpho-molecular features extracted from pre-treatment biopsies was evaluated. Starting with molecular predictors, no predictive features of expression levels of selected EMT-related markers (Catenin beta-1 & E-cadherin, supp. Fig. 29-32), including CDX2 (Fig. 5a & 5b) were found. For morphological features, neither *cluster size* (Fig. 5c & 5d) nor *interaction score* (supp. Fig. 33-34) can stratify patients into meaningful groups. Surprisingly, manual tumor bud counts (supp. Fig. 35-36) also did not significantly inform patient survival. This is potentially explained by the fact, that IF data leads to the over-estimation of TB, as stated by Aherne et al. [48]. However, our results from 2.5 show that EMT down-regulation in smaller clusters is a feature of tumor with infiltrative growth pattern. Therefore we suspected that combining molecular and morphological features can help with identifying tumor tissue undergoing active EMT processes. Therefore, a coordinated pattern of molecular and morphological features may be useful to indicate that the tumor tissue is undergoing active EMT processes. Indeed, when correlating CDX2 expression and *cluster size*, patients have a significantly lower survival probability for both OS (*p* < 0.0001) and DFS (*p* < 0.0001; Fig. 5e&f) if there is a strong correlation between the two features. This means that CDX2 down-regulation in small clusters is correlated with worse survival. To assess the independence of CDX2 correlation with *cluster size* as a predictor, cox proportional hazard (CoxPH) models were employed (see 4.6). In doing so, significance was retained for OS (*p* = 0.042; Fig. 5g) but not for DFS (*p* = 0.743; Fig. 5h). This can be explained by the tissue size per patient, making the data susceptible to variability in the sampling of tumor heterogeneity.

**Fig. 5.**
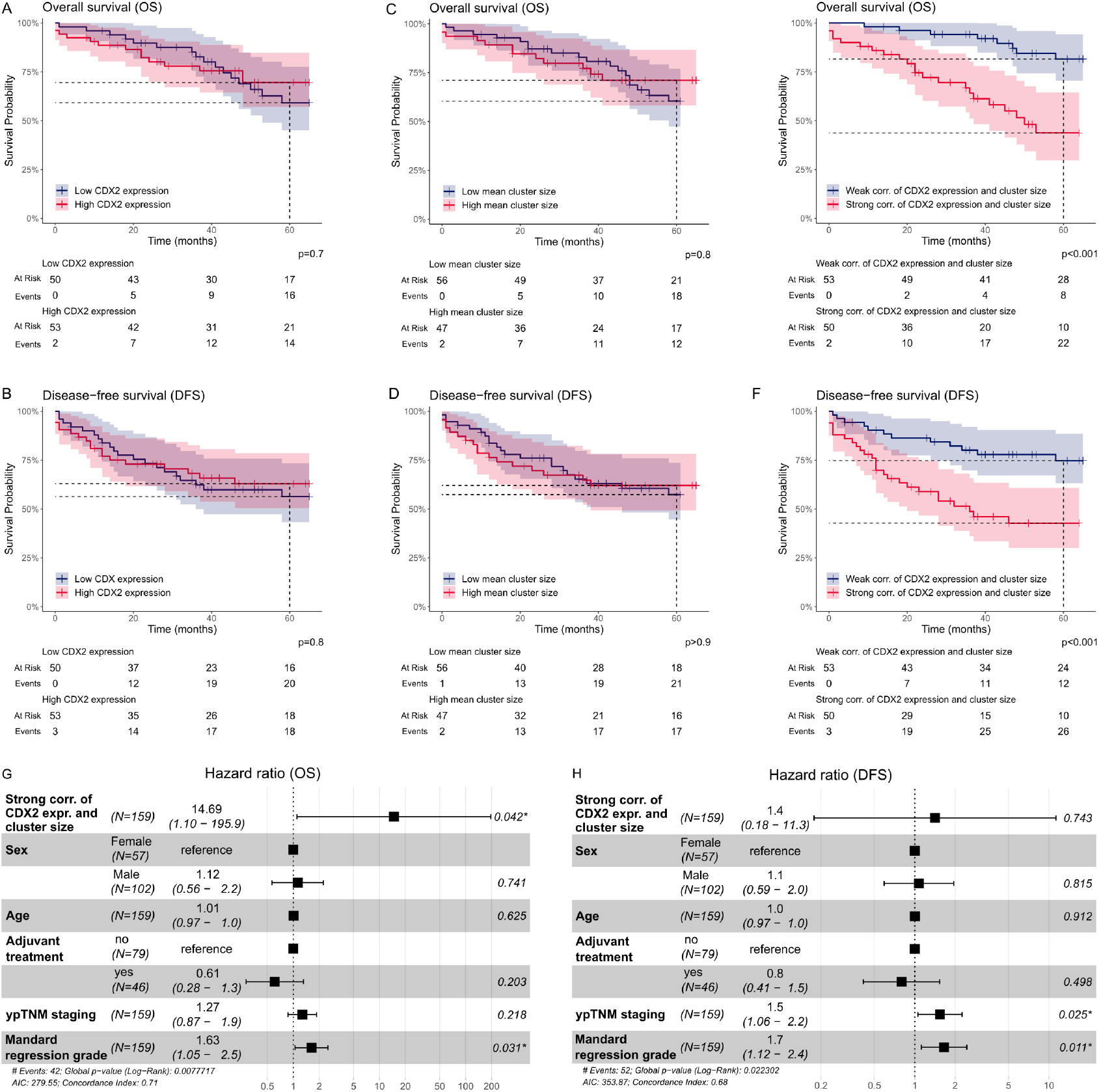
**(A)** Kaplan-Meier curves showing the overall survival of patients with high (red) or low (dark blue) CDX2 expression. **(B)** Disease-free survival of patients with high (red) or low (dark blue) CDX2 expression. **(C)** Overall survival of patients with high (red) or low (dark blue) mean *cluster size*. **(D)** Disease-free survival of patients with high (red) or low (dark blue) mean *cluster size*. **(E)** Overall survival of patients with low (red) or high (dark blue) CDX2 expression in small clusters. **(F)** Disease-free survival of patients with strong (red) or weak (dark blue) *cluster size* - CDX2 correlation. **(G)** Multivariate Cox-proportional hazards model for overall survival predicted by the correlation of CDX2 and *cluster size*. **(H)** Multivariate Cox-proportional hazards model for disease-free survival predicted by the correlation of CDX2 and *cluster size*.

Our new predictive feature was also compared with ypTNM staging and Dworak regression grading. When splitting the five grades of Dworak into three response groups first (Dworak 0 & 1 = “Bad response”; Dworak 2 = “Intermediate response”; Dworak 3 & 4 = “Good response”), no significance was found between CDX2 down-regulation in small clusters and either ypTNM (ANOVA; *p* = 0.064; supp. Fig. 45) or Dworak response groups (ANOVA; *p* = 0.249; supp. Fig. 46). CDX2 loss in smaller tumor clusters and tumor buds is therefore not able to predict Dworak regression grading. As all TRG methods rely on the fraction of tumor remnants and stromal tissue, the treatment effect on infiltratively growing tumors with an initially high fraction of stromal tissue is likely to be overestimated.

Continuing with survival analysis, log-rank tests and CoxPH analysis show a significant *EMT score* and *cluster size* correlation for OS (*p* = 0.034 and *p* = 0.047 respectively; supp. Fig. 37). This suggests that the larger EMT signature offers no benefit over CDX2 alone for survival prediction. CDX2 down-regulation along with an increased *interaction score* shows significance in Kaplan-Meier curves in both OS and DFS (*p* = 0.043 & *p* = 0.012 respectively; supp. Fig. 39-40) but not in CoxPH models (*p* = 0.41 & *p* = 0.63 respectively; supp. Fig. 43-44). Overall, we found association to worse survival in patients with CDX2 down-regulation in small clusters, suggesting that a loss of CDX2 in tumor buds and smaller clusters can be leveraged as an easily obtainable biomarker in biopsies.

## 3 Discussion

TB has been a widely investigated biomarker in colorectal cancer for the last three decades [19], [49], [50], [51], [52]. While its predictive power of metastatic progression in CRC was shown numerous times [19], [51], the biological justification remains opaque. There are likely two reasons for this: First, TB has originally been reported on 4‒5*µm*-thin sections [19] as a two-dimensional representation, therefore omitting the third dimension of the tumor. Recent studies have resolved TB in 3D, uncovering that although “real” tumor buds exist, many small 2D cell clusters are actually connected to larger tumor fragments, resulting in overestimation and incorrect biological interpretation of tumor bud processes in 2D [37], [53]. Second, while tumor buds are hypothesized to undergo EMT, this interaction is hard to demonstrate: In a clinical setting, the search for a highly reproducible biomarker for EMT is still ongoing [54], [55]. In an experimental setting, it is not possible to faithfully replicate the complex tumor microenvironment, which is essential for TB to emerge [56]. Therefore, to further our understanding of TB and its relation to EMT, we utilized sequential immunofluorescence (seqIF™) and computational image analysis methods.

In CRC, TB is recommended to be assessed at the invasive margin [20]. Comparing the two tumor locations *invasion front* and *tumor center*, we showed that TB had a bigger influence on marker expression than tumor location (Fig.1d-e). Furthermore, we can show differential epithelial protein expression of Catenin beta-1, CDX2 and E-cadherin between tumor buds and non-budding clusters within both tumor locations, suggesting that TB investigation within the tumor center is feasible (Fig. 1h). In fact, it was shown before that intratumoral budding is predictive of the occurrence of peritumoral budding [18], [25]. This also justifies the tumor bud investigation in the rectal biopsy cohort, as biopsies are routinely taken from the tumor center [23].

Applying our custom pipeline to the clinical rectal cancer cohort, we were able to replicate and extend on these results from the WSI and identified additional differentially expressed markers. In detail, we showed significant down-regulation of seven protein markers (Catenin beta-1, CDX2, E-cadherin, Ep-CAM, Lamin-B1, Ki-67, Caspase-3) in intratumoral buds of rectal biopsies. Differential Ki-67 and Ep-CAM expression levels in tumor buds were expected due to their previously described links to EMT [27], [57]. As Caspase-3 is a marker for cell apoptosis, lower levels in tumor buds can be explained by the absence necrotic regions often found within the larger tumor clusters [58]. Finally, Lamin-B1 down-regulation has been linked to EMT in non-small cell lung cancer [59] and is more broadly associated with cellular senescence [60]. However, Lamin-B1 down-regulation in tumor buds has not yet been shown in rectal cancer. Therefore, these findings warrant orthogonal validation. We therefore conclude that TB as a morphological feature can be evaluated on a TMA cohort, even though this feature is rare. This contrasts with a recent push in the field of spatial omics, where whole-slide analysis is preferred [37], as certain morphological features may be undersampled in TMA cores.

Exploring the principal components of tumor clusters in the rectal biopsy cohort, we found strong overlap in global expression patterns of tumor buds and non-budding clusters (Fig. 2c). To address the limitation of a binary classification step, we decided to introduce two alternative morphological features: *cluster size & interaction score* (described in sections 2.3, 2.4 & 4.5). We showed that both features not only correlate with an up-regulation of EMT (Fig. 2f & 3d) but also replicate the transition shown by EMT-related processes (Fig. 2e & 3b). We were able to reproduce *cluster size* results from Lin et al. [37] and showed the added value of our newly defined *interaction score*, as it also captures EMT in large clusters (Fig. 4d). Biologically, these results suggest both smaller tumor clusters and fibril-like structures being representative of the same EMT process (Fig. 4e). Such a similarity can be explained by the loss of information in 2D sections, as certain tumor clusters could be part of fibrils in a three-dimensional context. Leveraging the *cluster size* feature allows us to not only capture EMT at an advanced stage (represented by tumor buds) but also at its early stages. It is crucial to also quantify these transition states within biopsies, as they are more likely to be sampled than tumor buds in the tumor center, due to the rarity of intratumoral budding (ITB). Furthermore, these findings could add to the current definintion of Poorly Differentiated Clusters (PDCs) [35], [61], defined as clusters of more than four malignant cells without any visible lumen. We quantitatively found EMT upregulation up to a *cluster size* of 128, which could inform a size-based definition of PDCs [62]. This may open avenues towards re-considering the use of PDCs as a prognostic biomarker, both in colon cancer resections and rectal biopsies, as PDCs are more commonly found than ITB. It needs to be stated that our results show high variance, and therefore extracting a data-derived definition of PDCs is not possible with the data at hand but warrants further validation in additional cohorts.

We also found association of histological features and EMT within the colon cancer cohort. Differences of EMT upregulation are explained by tumor growth patterns: tumor border configuration has been suggested as a prognostic biomarker [36]. Strikingly, *cluster size* and *interaction score* had a stronger effect on EMT up-regulation in infiltrative tumors (Fig. 4c-d). Furthermore, infiltrative growth patterns correlated with CDX2-negativity in tumor buds (Fig. 4g). These findings suggest that active down-regulation of epithelial markers and specifically CDX2 in smaller tumor clusters is a way to capture infiltrative patterns and the aggressiveness of disease progression in static 2D patient samples.

We are aware of the limitations in the sample size of the colon cohort to make conclusions on interpatient heterogeneity, however, we can transfer these findings to the rectal biopsy cohort. When assessing various features on the second cohort, we concluded that none of the molecular or morphological features were able to predict survival on their own (Fig. 5a-d). Interestingly, TB counts do not correspond to patient survival either. As reported by Aherne et al. [48], counting TB visualized by immunofluorescence and PanCK expression can lead to an overestimation of tumor buds and lose the prognostic value. However, we do find significant interaction with OS and DFS when we inspect the active down-regulation of CDX2 in smaller clusters (Fig. 5e-g), showing that CDX2 could help identifying clinically relevant tumor buds. We also find significant log-rank tests for *EMT score* up-regulation in smaller tumor clusters, however, the same results cannot be shown in a CoxPH model. This can be explained by the arbitrary nature of the *EMT score* as well as the limitations of the cohort, as there is no possibility of assessing intra-tumor heterogeneity.

Further studies on larger cohorts with a more condensed panel, focusing on the TB markers E-cadherin, Catenin beta-1 and CDX2, will be needed to confirm these results and to assess the robustness of our biomarker. In the future, the search for novel TB protein markers needs to be focused on the discovery of up-regulated proteins: we did not find up-regulation of the proposed marker proteins Vimentin, ZEB1 and nuclear Catenin beta-1 in our data [30], [31], [32]. These findings can be explained by either an absence of expression or a non-optimal protocol for these markers within the panel. Furthermore, increasing the three-dimensional potential of the dataset with sequential cuts would add certainty to the validity of the extracted features.

In this work, we built a custom pipeline for sequential Immunofluorescence (seqIF™) to segment and classify tumor buds. We tackled the biological question of EMT and its link to tumor budding, and quantified morphological transition states. By quantifying *cluster size* and tumor-stroma interactions, we uncovered similar EMT processes in fibrils, as seen in tumor buds, and found active up-regulation of EMT in infiltratively growing cancers. This shows the importance of considering TB, fibril-like structures and larger tumor clusters as transitory states and not binary features. Finally, this study provides a novel dataset, consisting of a 31-plex immunofluorescence panel and 160 patients, the first hyperplex dataset of rectal cancer biopsies of such size. Leveraging the potential of this dataset, we show the relevance of CDX2 expression in small tumor clusters and its relation to survival, which may be used as a biomarker of infiltrative growth.

## 4 Methods

### 4.1 Cohort

The data used in this study consists of two cohorts. The first cohort consists of five whole slide images (WSI) of the colon topology from five patients (Fig. 6a I). The donor patients of this cohort did not receive any treatment. The tissue specimens were processed with the COMET™ protocol, with 24 markers being imaged sequentially (see 4.2). Each image is of physical size ∼9 × 9*mm* and has a resolution of 0.23 × 0.23*µm*, resulting in a digital size of ∼39580 × 39620*px*. As these images were part of the panel establishment, the number of visualized proteins range from 18 to 24 markers (see 4.2).

**Fig. 6.**
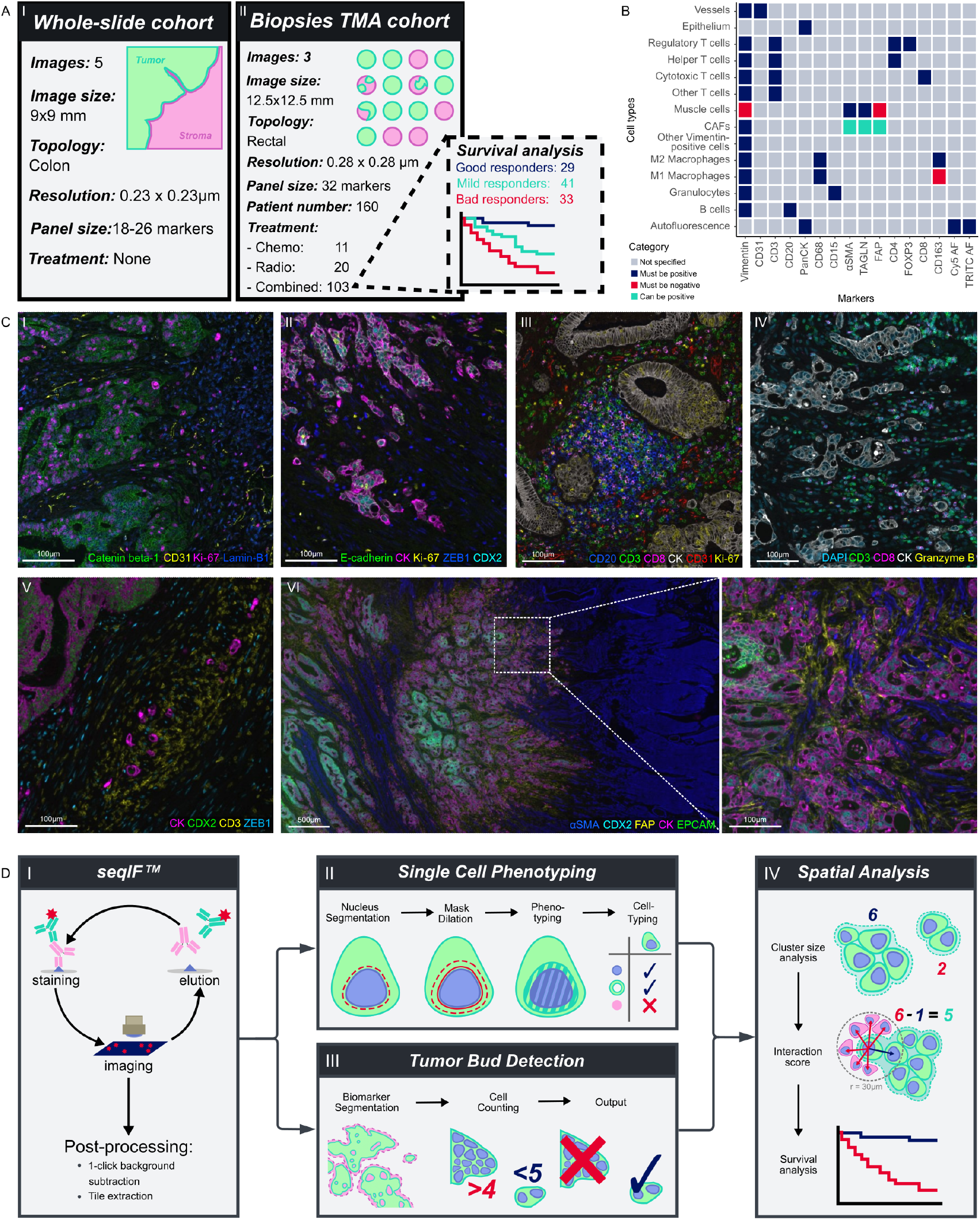
**(A)** Graphical scheme describing both available cohorts. **(B)** Cell typing instructions, which defines which markers are known to be positive (or negative) for the cell types of interest. **(C)** Composite images of the complete 31-plex images, grouped by functional classes: markers involved in EMT (I & II), immune cells (III), markers for T-cell characterization (IV), markers for fibroblast characterization (V). **(D)** Schematic of the custom seqIF™ pipeline.

The second cohort consists of three tissue micro-arrays (TMA) with a total of 160 locally-advanced rectal cancer (LARC) patients (Fig. 6a II), where each TMA core (∅= 1*mm*) corresponds to one patient. Each TMA also carries three control cores (normal colon epithelium, tonsil epithelium & tonsil germinal center). The three images were scanned with a resolution of 0.28 × 0.28*µm* and a physical size of 12.5 × 12.5*mm*. A total of 31 different epitopes were visualized on each slide (see 4.2). 103 patients received radio-chemotherapy, 20 patients received only radio-therapy and 11 patients received only chemotherapy. The treatment regimen is unknown for 26 patients.

### 4.2 Immunofluorescence

Sequential immunofluorescence (seqIF™) protocol was performed using the COMET™ platform for the spatial detection of 31 markers. The OME-TIFF output contains images of DAPI for nuclear staining, two autofluorescence channels for background subtraction, and 31 single marker channels which were used for downstream image analysis. Example composite images are shown in Fig. 6c, and single-channel images are given in supplementary figure 47.

Automated seqIF™ staining, imaging and antibody elution was performed on the COMET™ platform (Lunaphore Technologies). Slides underwent 20 cycles of iterative staining and imaging, followed by elution of the primary and secondary antibodies [41].

In brief, FFPE slides were preprocessed with PT Module (Epredia) with Dewax and HIER Buffer H (TA999-DHBH, Epredia) for 60 min at 102°C. Subsequently, slides were rinsed and stored in Multistaining Buffer (BU06, Lunaphore Technologies) until use.

The protocol was established as 24plex and expanded to 31plex for the rectal biopsy cohort. The protocol template was generated using the COMET™ Control Software, and reagents were loaded onto the instrument to perform the seqIF™ protocol. List of primary antibodies is enclosed in supplementary table 2. Secondary antibodies were used as a mix of two and DAPI: Alexa Fluor Plus 647 goat anti-mouse (Lunaphore Technologies, cat no: DR647MS, 1/200 dilution), Alexa Fluor Plus 555 goat anti-rabbit (Lunaphore Technologies, cat no: DR555RB, 1/100 dilution), Alexa Fluor Plus 647 goat anti-rabbit (Lunaphore Technologies, cat no: DR647RB, 1/200 dilution), Alexa Fluor Plus 555 goat anti-mouse (Lunaphore Technologies, cat no: DR555RB, 1/100 dilution). Nuclear signal was detected with DAPI (Lunaphore Technologies, DR100) by dynamic incubation of 2 min. All primary antibodies were diluted in Multistaining Buffer (BU06, Lunaphore Technologies) and secondary antibody mixes were diluted in Intercept T20 (TBS) Antibody Diluent (Licorbio). All the primary antibodies were incubated for 8 min and all the secondary antibody mixes were incubated for 2min. Images were taken with default exposure times in COMET™. Elution step lasted 2 min for each cycle and was performed with Elution Buffer (BU07-L, Lunaphore Technologies) at 37°C. Quenching step lasted for 30 sec and was performed with Quenching Buffer (BU08-L, Lunaphore Technologies). Imaging step was performed with Imaging Buffer (BU09, Lunaphore Technologies). The seqIF™ protocol in COMET™ resulted in a multi-layer OME-TIFF file where the imaging outputs from each cycle are stitched and aligned. COMET™ OME-TIFF output contains DAPI image, intrinsic tissue autofluorescence in TRITC and Cy5 channels, a single fluorescent layer per marker and single layer per additional image post-elution.

### 4.3 Pipeline

#### 4.3.1 Image preprocessing

Each COMET™ run produces multiple 3-channel images, consisting of one DAPI-channel plus two administered antibody pairs. Registration of these images is done via the DAPI-channel of each channel and happens within the COMET™ software. The images are supplied in the OME.TIFF format. Tissue autofluorescence is subtracted using the “HORIZON™ Viewer” software (Fig. 6d I). Whole-slide images are split into tiles of size 1921 × 1925*px*, resulting in 420 tiles per image. Each tile has an overlap of 40*px* in both dimensions. TMA images are first de-arrayed using QuPath [74]. Furthermore, each de-arrayed core is split into 4 tiles of size 2420 × 2420*px*. The tiles have an overlap of 30*px* in both dimensions.

#### 4.3.2 Cell phenotyping

Nucleus segmentation is performed on the DAPI-channel using StarDist [75] (Fig. 6d II). Each tile’s DAPI expression is first normalized before applying the StarDist pipeline. The resulting segmentation is dilated by a factor of 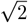 in both axes. To prohibit segmentation overlap, watershed is applied to assign each pixel to one single cell instance. Per cell, marker expression is measured and averaged within the nuclear and cytoplasmic compartment. Each marker is *log*10-transformed and subsequently normalized per slide using quantile normalization

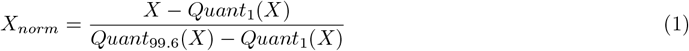

where *Quant*_1_(*X*) and *Quant*_99.6_(*X*) correspond to the 1^st^ and 99.6^th^ quantile of each marker for each slide. The transformed and normalized data is then converted to the `anndata` format [76], [77] for downstream operations.

#### 4.3.3 Cell typing

A set of instructions is given to represent well-defined cell types (Fig. 6b) such as epithelium, vessels, T-cells and their various subtypes, muscle cells and more. Cell types such as “other Vimentin-positive cells” and “Other T cells” are used to catch cells that cannot be further subtyped, as no other distinguishing markers are expressed. These cells are subsequently removed from the analysis. Furthermore, two autofluorescence channels (TRITC & Cy5) are used to find epithelial cells with strong autofluorescent signals, which in turn are removed from downstream analysis. This is used to filter out false-positive erythrocytes and necrotic regions. Before applying the set of instructions to the expression data, each marker expression distribution is rescaled by fitting a gaussian mixture model (GMM). The data is then rescaled such that the intercept of the two fitted gaussian distribution is set to 0.5, and that all values range between 0 and 1. A cell is considered positive for a marker if its marker expression value exceed 0.5. ”” Finally, the set of instructions is applied to the thresholded expression patterns of each cell. In the special case of cancer-associated fibroblasts (CAFs), at least one marker of FAP, *α*SMA or TAGLN needed to be expressed (Fig. 6b). Furthermore, to differentiate CAFs from muscle cells, only Vimentin-negative cells were classified as muscle cells. Cell typing workflow was written in `python` using the `scanpy` [78] and `scimap` [79] packages.

### 4.4 Tumor bud detection

To segment epithelial tissue, a random forest segmentation method was applied (Fig. 6d III). For each image of both cohorts, a random forest [80] was trained on 5 tiles per image. Scribble annotation [81] was given for both the background and the epithelium class. The trees were split on nine pixel features per channel, based on intensity (three per channel) and texture (six per channel). To capture both small-scale and large scale features, gaussian blur (*σ* = {1, 2.45, 6}) was applied. Retraining the pixel classification algorithm for each image was conducted to account for inter-image differences (e.g. resolution, average marker intensity, image-specific artifacts) For each training tile, 10 trees of a resulting forest are trained, resulting in a 50-tree forest. The generated tissue masks are post-processed by removing small holes followed by applied mask erosion and removal of small objects. Taking into account the nuclei segmentation and cell typing results, the number of cells within each uniquely segmented tumor area is assessed (Fig. 2d). Given the definition of tumor buds [20], a threshold of *n* < 5 connected malignant epithelial cells is applied to classify tumor buds and non-budding clusters (Fig. 6d III). Random forest segmentation was applied using the `python` package `skimage` [82], subsequent mask operations were done using packages `skimage` and `scipy` [83].

### 4.5 Post processing

Several post-processing steps were applied to the results. Within the context of the colon cancer cohort, leiden clustering [84] identified artifact clusters (necrosis, autofluorescent tissue) and normal mucosa. These clusters were subsequently removed from downstream analysis [85]. One image also showed “funky stroma” (PanCK-positive stroma [30]), which was removed from the analysis with annotations created in QuPath [74]. Furthermore, substantial batch-effects between the images were corrected using ComBat [86] implemented in scanpy [78]. Finally, *EMT scores* and *interaction scores* are calculated. Markers CDX2, Catenin beta-1 & E-cadherin were averaged and trimmed off at 0 and 1 to create the *EMT-score*. To calculate the *interaction score* (Fig. 6d IV), the number of cancer-associated fibroblasts (CAFs) within 30*µm* of each epithelial cell were quantified to find the degree of stromal interaction. We excluded immune cells from the count, as this should not be a quantification of immune reaction. Cell types have been defined using the previously described set of instructions (see 4.3.3). Furthermore, epithelial interaction was captured by counting the number of other malignant cells within 30*µm* of each tumor cell (Fig. 3a). Subtracting the epithelial interaction from the stromal interaction to get the *interaction score*, values above 0 will correspond to stronger stromal interaction and values below 0 correspond to stronger epithelial interaction. Furthermore, the *interaction score* was used for artifact filtering. Small tissue areas that are not connected to any stromal tissue are likely to have emerged from either cutting or punching artifacts, or represent pseudobuds [47]. Therefore, tumor buds with a mean *interaction score* of 0 or lower are removed from downstream analyses. PCA and UMAP embeddings were created using the `python` package `scanpy` [78].

### 4.6 Statistical analysis

Wilcoxon tests were conducted to assess differential expression between tumor locations (Fig. 1b-d). Tumor locations were annotated manually: the invasion front was defined as a 500*µm* radius zone around the deepest invasion. The tumor center is defined as the remaining cancer tissue (Fig. 1b). Tissue fragments without apparent invasion front were excluded from tumor location assessment. Paired wilcoxon tests have been conducted to assess differential expression between tumor buds and larger tumor clusters within the same tile or core (Fig. 1e & h). All differential expression tests were Bonferroni-corrected. All statistical tests were conducted in `R`. Survival analyses were conducted with log-rank tests [87], using disease-free survival and overall survival (as defined by Punt et al. [88]) as endpoints. Patients that did not receive nCRT were excluded from survival analysis. The independence of the predictors from known prognostic markers such as adjuvant treatment [89], ypTNM staging [13] and Mandard regression grading [12] was assessed using a Cox proportional hazard model [90]. Survival analysis was conducted using `R` packages `survival` [91], `survminer` [92] and `ggsurvfit` [93]. All figures were created using `ggplot2` [94].

## 5 Acknowledgements

M.G. was supported by the Swiss cancer league under grant number KFS-5786-02-2023-R. M.G. and C.S.D. were supported by Innosuisse through innovation project 51906.1 IP-LS.

## 6 Data availability

Five whole slide images from the colon cancer cohort will be made available upon publication of the paper in the corresponding repository: XX-XXX-XX

## 7 Code availability

All code used in this study is accessible on github: https://github.com/digitalpathologybern/MxIF-pipeline

## 8 Authors’ contribution

- **Conception and design:** M.G., C.S.D, I.Z.
- **Development of methodology:** M.G.
- **Acquisition of data:** C.S.D
- **Analysis and interpretation of data (e.g**., **statistical analysis, biostatistics, computational analysis):** M.G.
- **Writing, review, and/or revision of the manuscript:**M.G., C.S.D., H.L.W., A.L., C.G.M., J.K, A.K., P.K., T.K., M.D.B., M.W., I.Z.
- **Administrative, technical, or material support (i.e**., **reporting or organizing data, constructing**
- **databases):** M.G., C.S.D, C.G.M., M.D.B. H.L.W., I.Z.
- **Study supervision:** M.W., I.Z.

