## Supplemental figures for "Morpho-molecular features of Epithelial Mesenchymal Transition associate with clinical outcome in patients with rectal cancer"

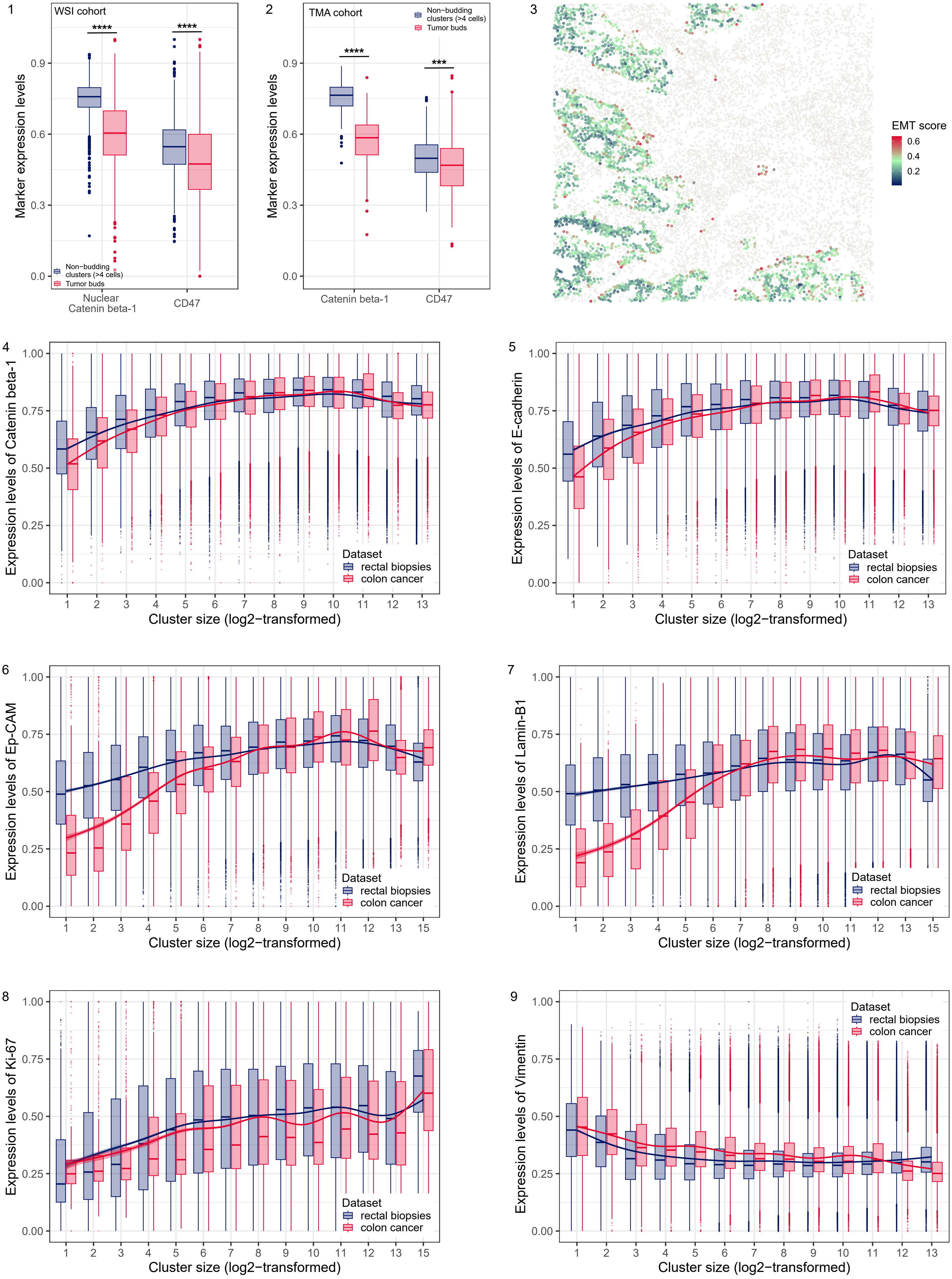
1
WSI cohort
Marker expression levels
Non−budding clusters (>4 cells)
Tumor buds
Nuclear Catenin beta-1
CD47
2
TMA cohort
Non−budding clusters (>4 cells)
Tumor buds
Marker expression levels
Catenin beta-1
CD47
3
EMT score
0.6
0.4
0.2
4
Expression levels of Catenin beta-1
Dataset
rectal biopsies
colon cancer
Cluster size (log2−transformed)
5
Expression levels of E-cadherin
Dataset
rectal biopsies
colon cancer
Cluster size (log2−transformed)
6
Expression levels of Ep-CAM
Dataset
rectal biopsies
colon cancer
Cluster size (log2−transformed)
7
Expression levels of Lamin-B1
Dataset
rectal biopsies
colon cancer
Cluster size (log2−transformed)
8
Expression levels of Ki-67
Dataset
rectal biopsies
colon cancer
Cluster size (log2−transformed)
9
Expression levels of Vimentin
Dataset
rectal biopsies
colon cancer
Cluster size (log2−transformed)

10

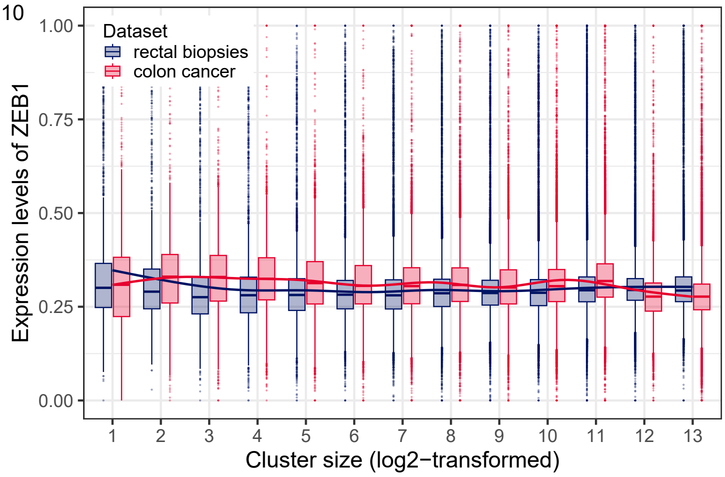


11

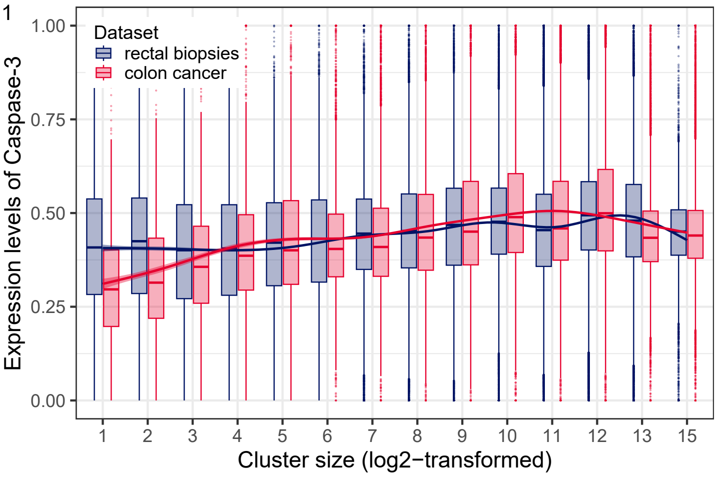


12

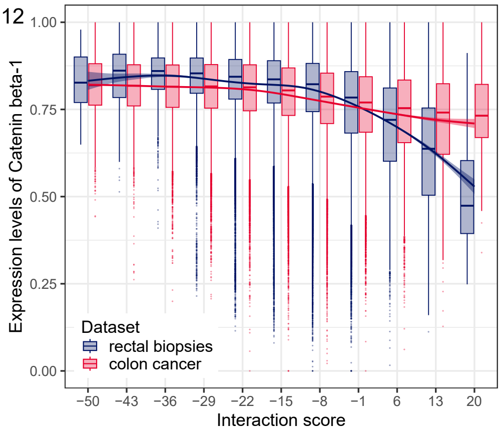


13

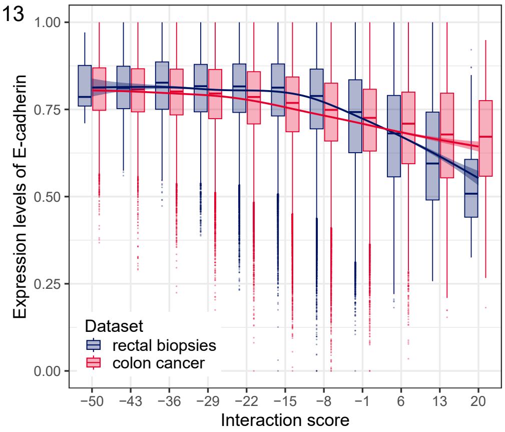


14

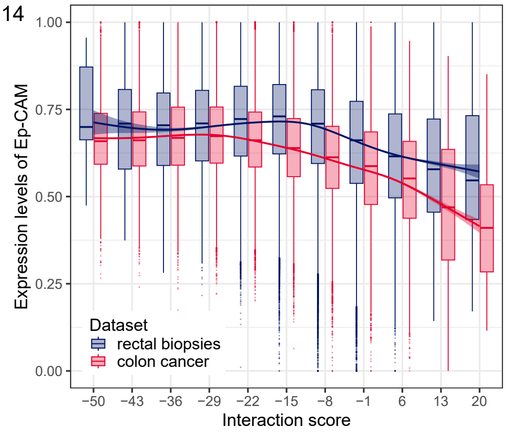


15

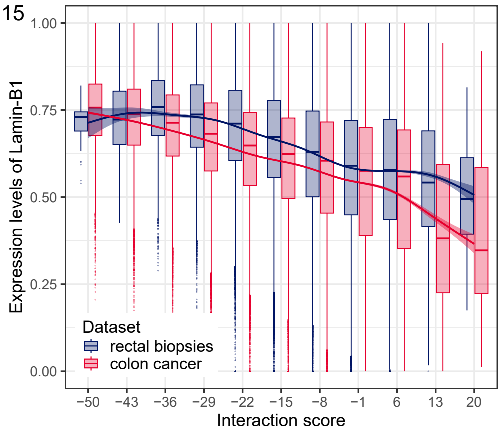


16

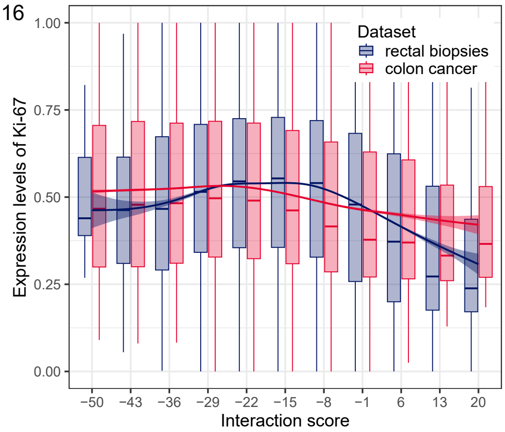


17

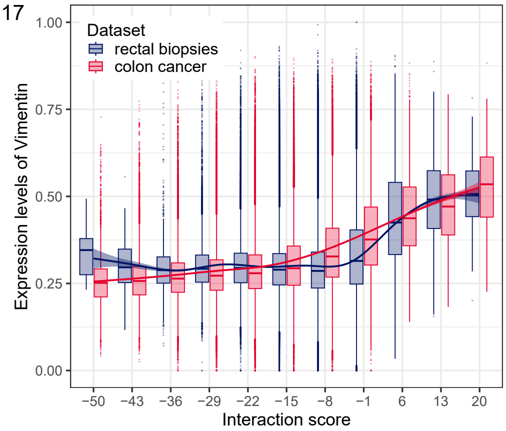


18

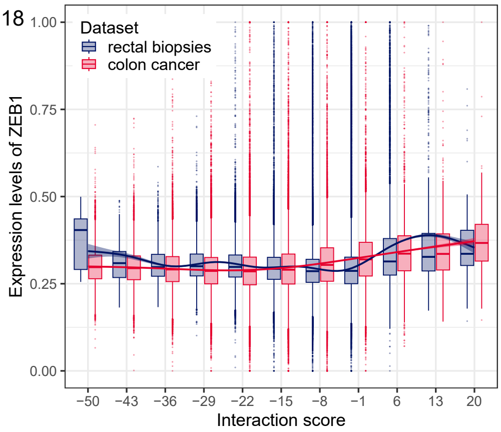


19

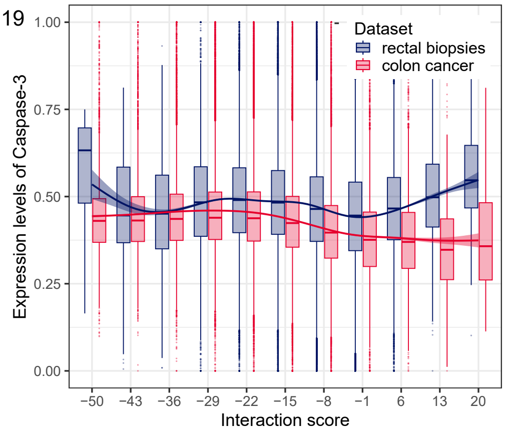


20

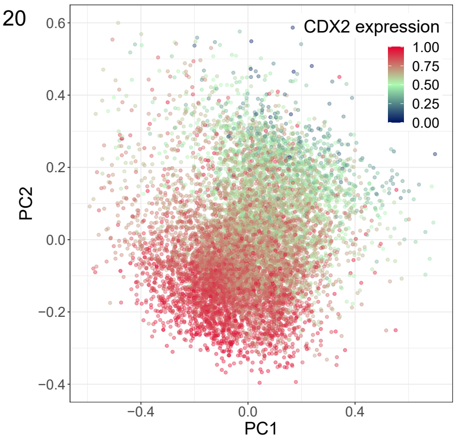

21

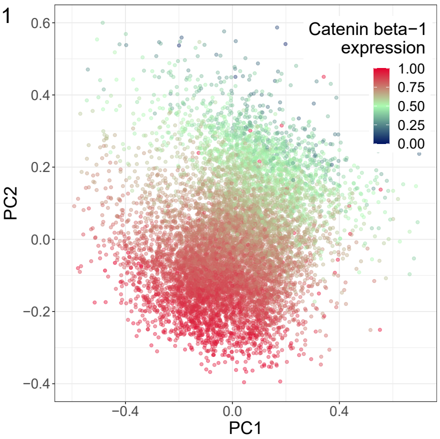


22

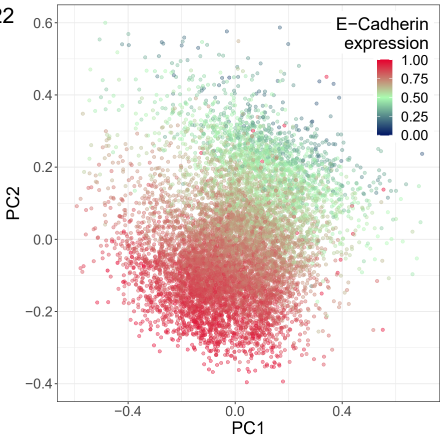


23

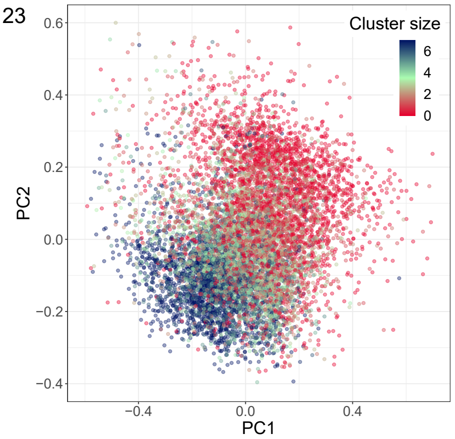


24

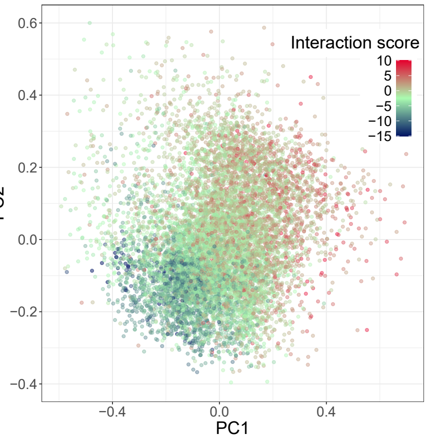


25

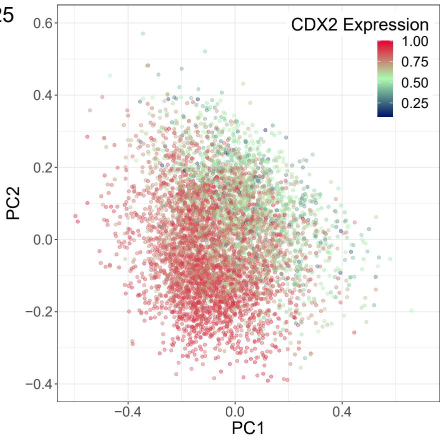


26

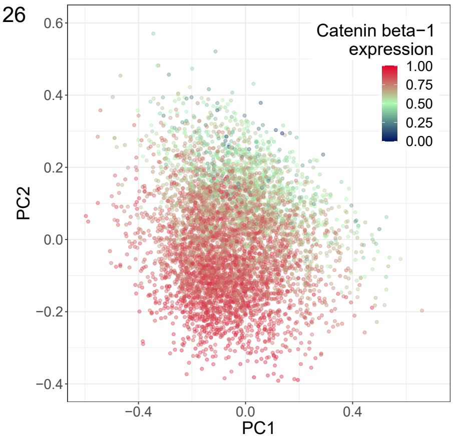


27

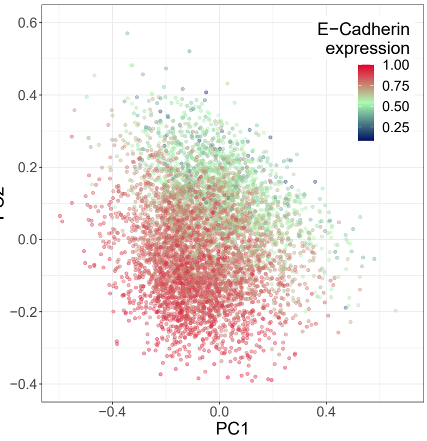


28

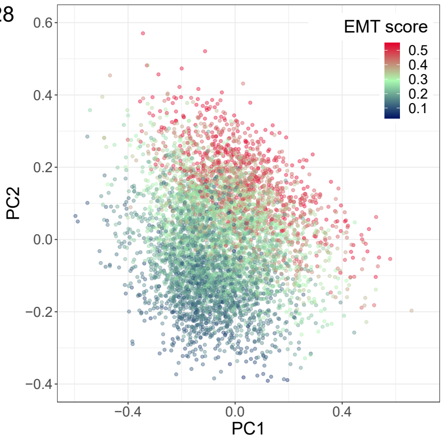


29

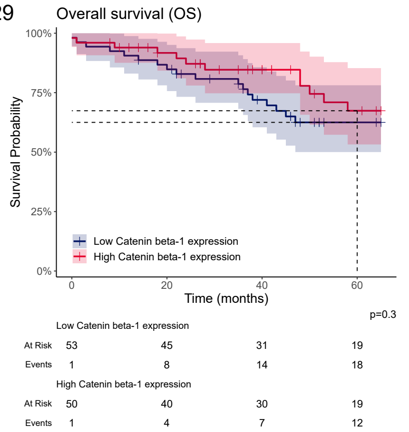


30

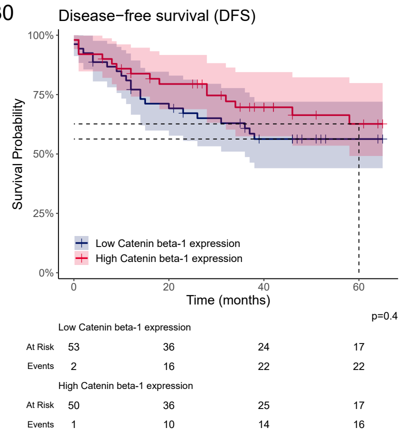


31

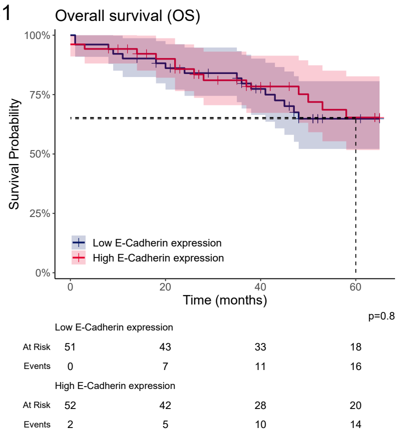


32

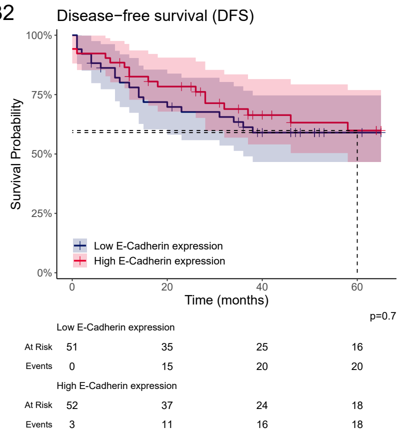

33

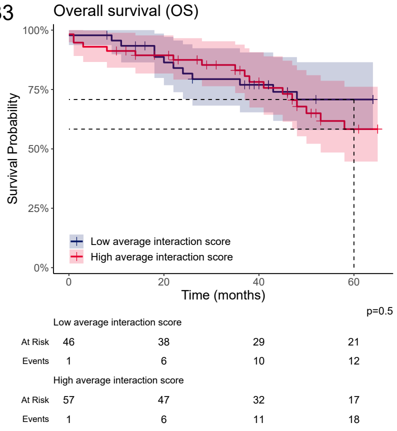


34

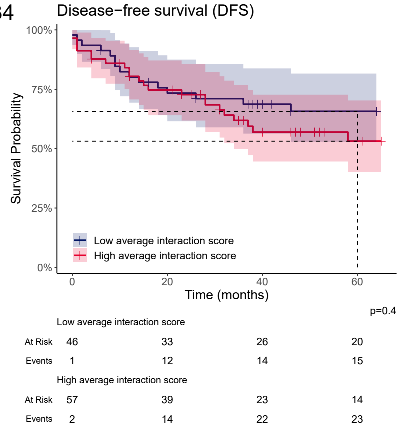


35

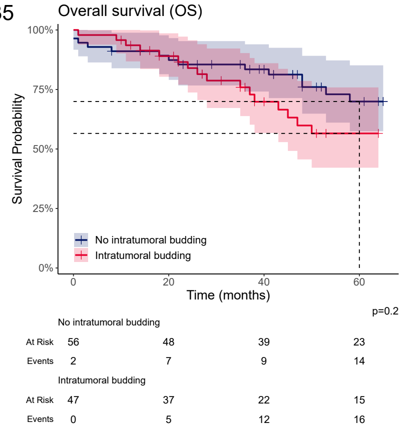


36

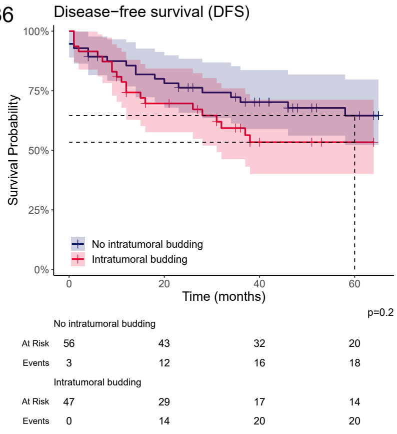


37

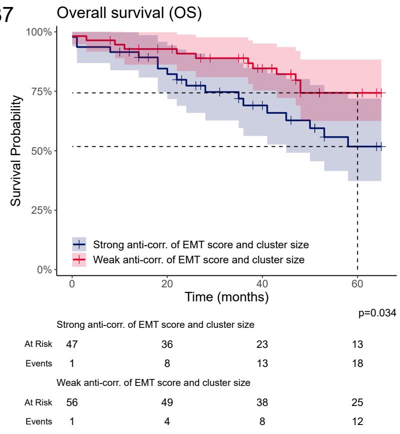


38

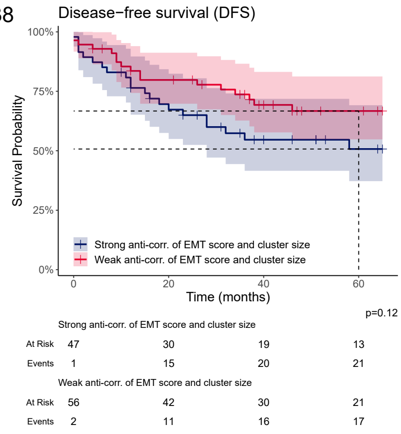


39

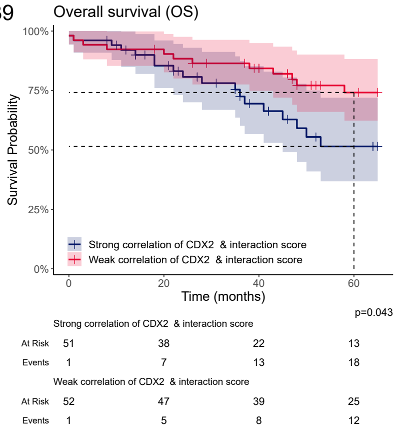


40

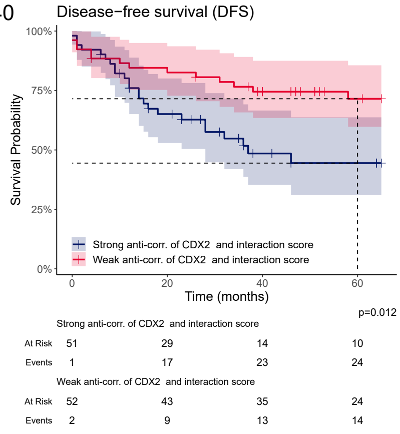


41

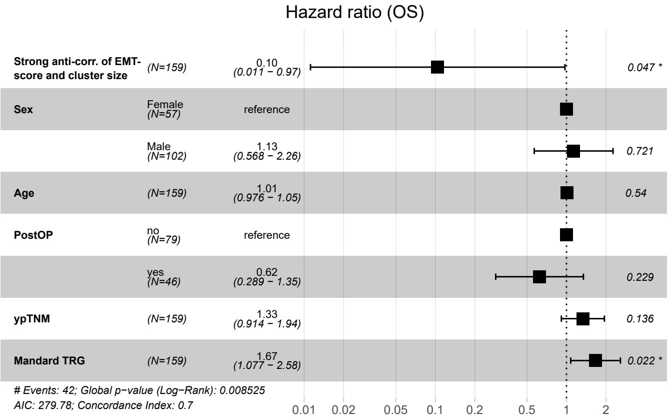


42

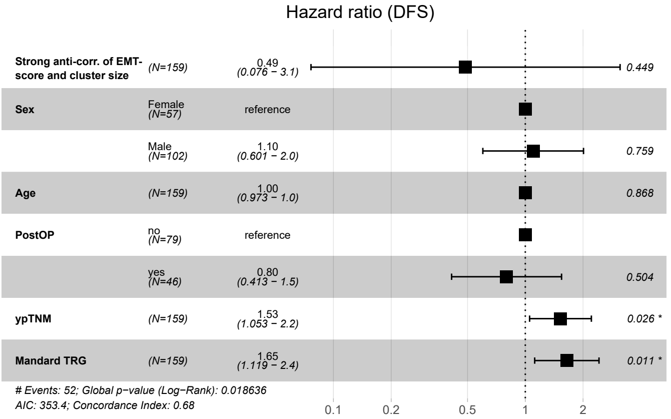

43

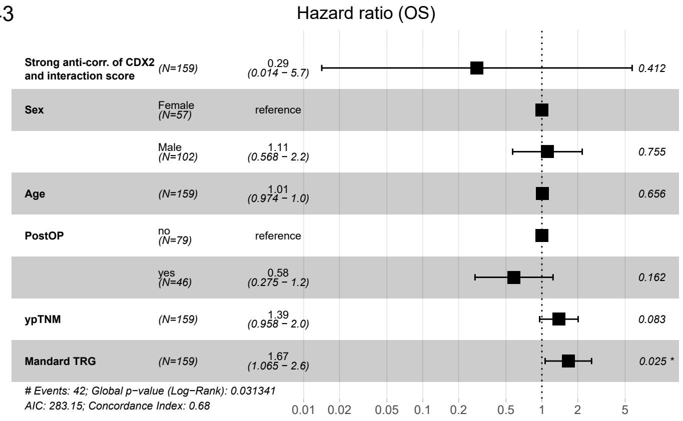

Hazard ratio (OS)
Strong anti-corr. of CDX2 and interaction score (N=159) 0.29 (0.014 – 5.7) 0.412
Sex Female (N=57) reference
Male (N=102) 1.11 (0.568 – 2.2) 0.755
Age (N=159) 1.01 (0.974 – 1.0) 0.656
PostOP no (N=79) reference
yes (N=46) 0.58 (0.275 – 1.2) 0.162
ypTNM (N=159) 1.39 (0.958 – 2.0) 0.083
Mandard TRG (N=159) 1.67 (1.065 – 2.6) 0.025 *
### Events: 42; Global p-value (Log-Rank): 0.031341
AIC: 283.15; Concordance Index: 0.68
0.01 0.02 0.05 0.1 0.2 0.5 1 2 5


44

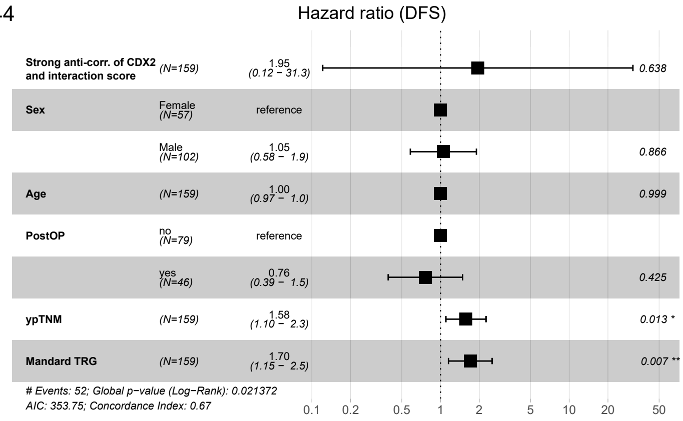

Hazard ratio (DFS)
Strong anti-corr. of CDX2 and interaction score (N=159) 1.95 (0.12 – 31.3) 0.638
Sex Female (N=57) reference
Male (N=102) 1.05 (0.58 – 1.9) 0.866
Age (N=159) 1.00 (0.97 – 1.0) 0.999
PostOP no (N=79) reference
yes (N=46) 0.76 (0.39 – 1.5) 0.425
ypTNM (N=159) 1.58 (1.10 – 2.3) 0.013 *
Mandard TRG (N=159) 1.70 (1.15 – 2.5) 0.007 **
### Events: 52; Global p-value (Log-Rank): 0.021372
AIC: 353.75; Concordance Index: 0.67
0.1 0.2 0.5 1 2 5 10 20 50


45

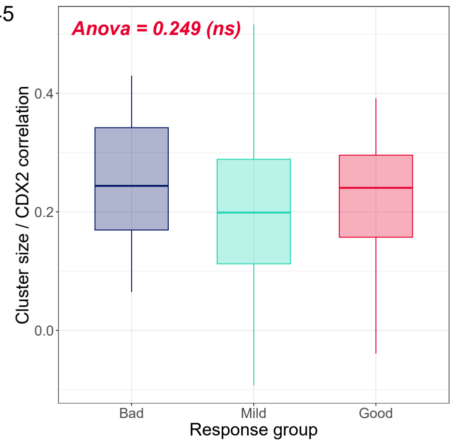

Anova = 0.249 (ns)
Cluster size / CDX2 correlation
0.4
0.2
0.0
Bad
Mild
Good
Response group


46

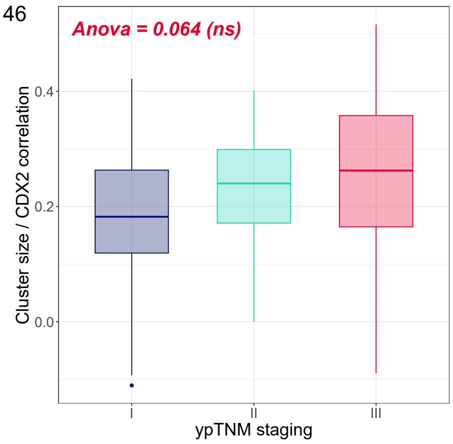

Anova = 0.064 (ns)
Cluster size / CDX2 correlation
0.4
0.2
0.0
I
II
III
ypTNM staging

47

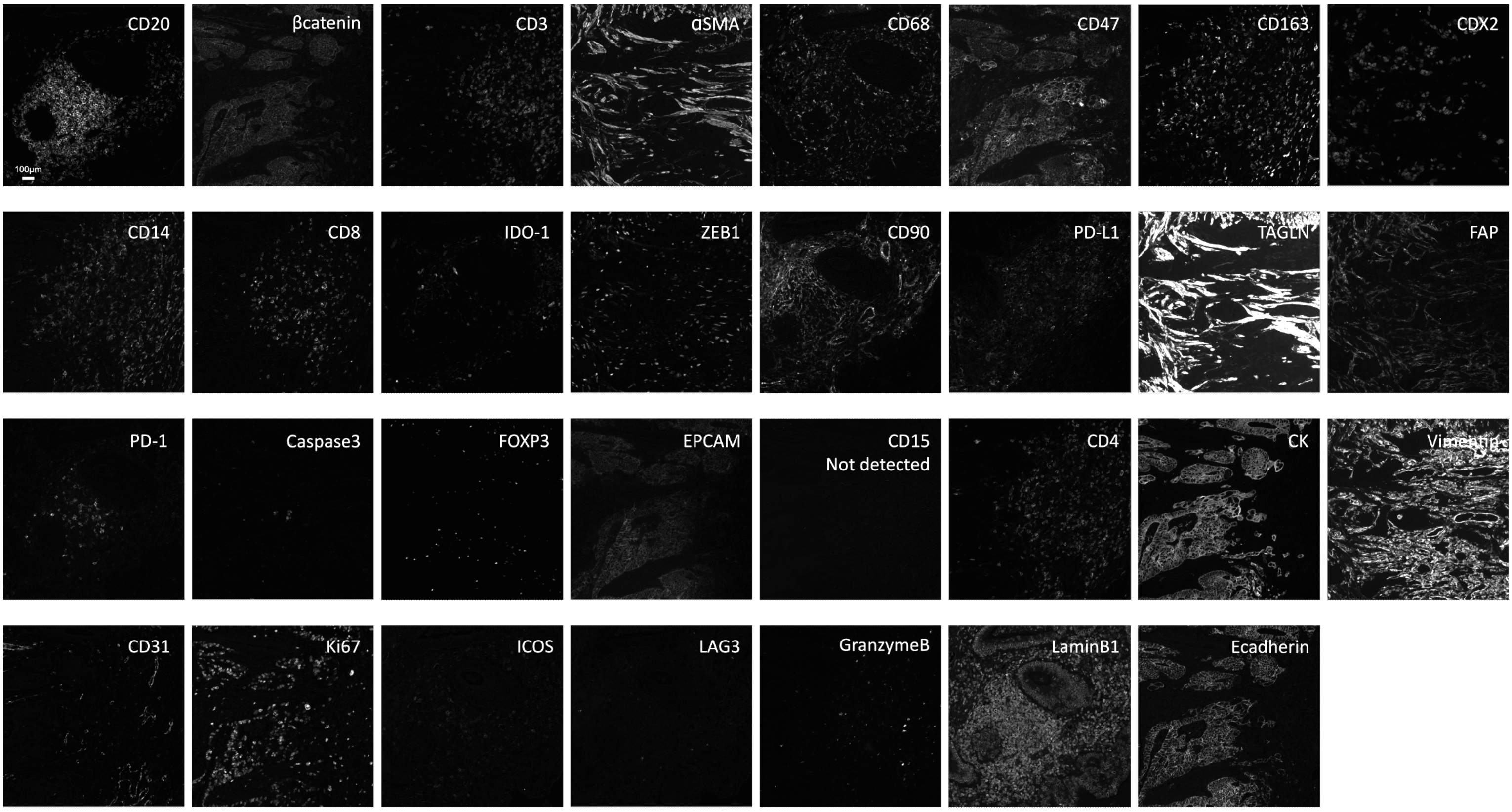
CD20
βcatenin
CD3
αSMA
CD68
CD47
CD163
CDX2
100µm
CD14
CD8
IDO-1
ZEB1
CD90
PD-L1
TAGLN
FAP
PD-1
Caspase3
FOXP3
EPCAM
CD15
Not detected
CD4
CK
Vimentin
CD31
Ki67
ICOS
LAG3
GranzymeB
LaminB1
Ecadherin

48

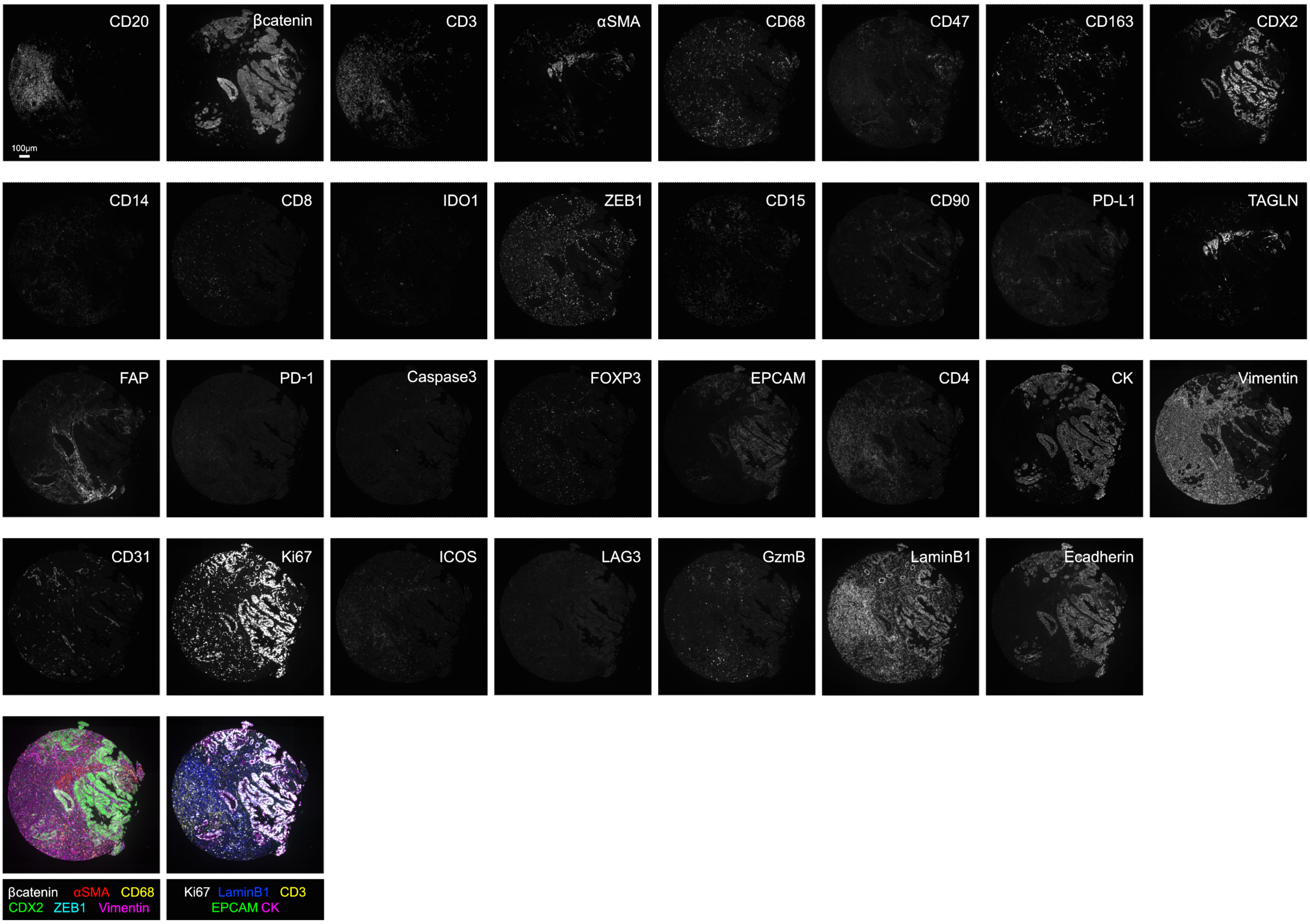
CD20
βcatenin
CD3
αSMA
CD68
CD47
CD163
CDX2
100µm
CD14
CD8
IDO1
ZEB1
CD15
CD90
PD-L1
TAGLN
FAP
PD-1
Caspase3
FOXP3
EPCAM
CD4
CK
Vimentin
CD31
Ki67
ICOS
LAG3
GzmB
LaminB1
Ecadherin
βcatenin αSMA CD68
CDX2 ZEB1 Vimentin
Ki67 LaminB1 CD3
EPCAM CK

**Supplementary figures: (1-2)** Differential expression of nuclear Catenin beta-1 and CD47 in both the colon cancer and rectal biopsy cohort . (3) Visualization of the EMT score in the same tumor region as figure 1A. **(4-11)** Correlation of log2-transformed *cluster size* and markers Catenin beta-1, E-cadherin, Ep-CAM, Lamin-B1, Ki-67, ZEB1, Vimentin, and Caspase-3. **(12-19)** Correlation of *interaction score* and markers Catenin beta-1, E-cadherin, Ep-CAM, Lamin-B1, Ki-67, ZEB1, Vimentin, and Caspase-3. **(20-24)** PCA visualizations of every tumor cluster in the colon cancer cohort. Colors signify CDX2 expression, Catenin beta-1 expression, E-cadherin expression, *cluster size* and *interaction score*. **(25-28)** PCA visualizations of every tumor cluster in the rectal biopsy cohort. Colors signify CDX2 expression, Catenin beta-1 expression, E-cadherin expression and *EMT score*. **(29-40)** Kaplan-meier curves with the predictors “Catenin beta-1 expression”, “E-cadherin expression”, “*interaction score*”, “Intratumoral budding”, “EMT score in smaller clusters” and “Strong correlation of CDX2 & interaction score”. **(41-44)** Cox-proportional Hazard models for predictors “EMT score in smaller clusters” and “Strong correlation of CDX2 & interaction score”. **(45)** Boxplots showing the strength of correlation between CDX2 and log2-transformed *cluster size* for the different ypTNM stages. Stage 4 was excluded due to the low amount of cases. **(46)** Boxplots showing the strength of correlation between CDX2 and log2-transformed *cluster size* for the different Response groups. **(47)** Single-channel images of the complete 31-plex panel. **(48)** Single-channel images of the complete 31-plex panel are shown on a representative rectal biopsy core, together with two composite examples.
